# Ribosome biogenesis bottlenecks reveal vulnerabilities in cancer

**DOI:** 10.64898/2026.04.20.719509

**Authors:** Lifei Jiang, Qiwei Yu, Sofia A. Quinodoz, Jordy F. Botello, Sabreen Alam, Jing Xia, Jonida Trako, Troy J. Comi, Aya A. Abu-Alfa, Yong Wei, Andrej Košmrlj, Yibin Kang, Clifford P. Brangwynne

## Abstract

Cell growth requires elevated protein synthesis, which depends on the production of ribosomes. Ribosome biogenesis is a complex, multi-step pathway in which newly transcribed precursor ribosomal RNA (rRNA) undergoes coordinated processing and assembly in the nucleolus to produce the small and large ribosomal subunits (SSU and LSU).^1–3^ Oncogene activation stimulates rRNA transcription and processing, giving rise to enlarged nucleoli that produce thousands of ribosomes every minute.^4,5^ However, efficient ribosome production requires tight coordination across numerous maturation steps, and it remains unclear if elevated rDNA transcription is proportionally converted into mature ribosomes, or whether imperfect coordination constrains the output yield. Here, we quantify pre-rRNA transcription (input) and compare it with newly-assembled cytoplasmic ribosomes (output), revealing that oncogene activation reduces the efficiency of ribosome production. Using a quantitative pulse-chase sequencing approach with mathematical modeling to resolve rRNA maturation kinetics, we found that oncogene activation creates late-stage processing bottlenecks, characterized by delayed precursor maturation and increased degradation. Perturbation of late-stage ribosome biogenesis factors preferentially impaired oncogene-driven cell growth, and limited tumor growth in mouse models, suggesting that these bottlenecks represent selective vulnerabilities in cancer, created by imbalanced biosynthetic flux. Together, these findings reveal that oncogene-driven ribosome production is imperfectly coordinated across maturation steps, and suggest that capacity limits in multi-step assembly pathways may be therapeutically exploitable in cancer and other diseases.

## Introduction

Ribosomes are essential molecular machines that translate messenger RNAs into proteins. Ribosome biogenesis requires coordinated production of rRNA and ribosomal proteins to produce up to thousands of ribosomes per minute^2,3,6,7^, representing one of the most energetically costly processes in proliferating cells.^8,9^ This multistep maturation pathway is orchestrated by ∼200 processing and assembly factors, underscoring the remarkable complexity of ribosome production.^2^ To coordinate these reactions, ribosome biogenesis is spatially organized across the cell. Within the nucleolus, pre-rRNAs are transcribed and subsequently cleaved, modified, and assembled with ribosomal proteins to form precursors of the small and large ribosomal subunits.^1,2,7^ These pre-ribosomal particles then undergo further processing in the nucleoplasm and are exported to the cytoplasm, where final maturation produces functional ribosomes.^10–12^

To sustain the high protein synthesis demands of rapid growth and proliferation, cancer cells markedly increase ribosome production by engaging oncogenic programs that stimulate ribosome biogenesis through multiple mechanisms.^13,14^ A key example is MYC-driven activation of RNA polymerase I (Pol I) transcription at ribosomal DNA (rDNA), through coordinated regulation of Pol I machinery that increases rRNA synthesis.^5,15^ Upstream oncogenic signals such as RAS further converge on MYC-dependent transcriptional programs that promote ribosome production and cell growth.^16,17^ Consistent with this upregulation of ribosome biogenesis, aberrant nucleolar morphology, often manifesting as nucleolar enlargement, has long served as a prognostic marker in cancer, correlating with increased tumor aggressiveness.^4^

Upregulation of ribosome biogenesis requires coordination across the entire assembly line, from rRNA transcription to processing and assembly with the increasing influx of ribosomal proteins. Newly transcribed pre-rRNAs must progress through an ordered cascade of thousands of reactions, including endonucleolytic cleavages, snoRNA-guided modifications, exonucleolytic trimming, and ribosomal protein assembly.^1,18,19^ Cleavage, modification, and trimming occur at multiple defined sites and are coupled to large-scale remodeling of pre-ribosomal particles, whereas aberrant or stalled intermediates are eliminated by nuclear surveillance pathways.^20–22^ Because ribosomal proteins are synthesized in the cytoplasm and imported into the nucleus, productive assembly further depends on matching the timing and stoichiometry of ribosomal protein availability to rRNA maturation across cellular compartments.^23^ MYC amplifies rRNA transcription while simultaneously increasing expression of ribosomal proteins and biogenesis factors,^24^ sharply elevating flux into the pathway. This raises the possibility that ribosome biogenesis operates as a capacity-limited assembly pathway, in which increased transcriptional input may not be proportionally converted into mature ribosome output if downstream processing steps become saturated.

Examining the degree to which these processes are quantitatively coordinated along the maturation pathway has been hampered by the lack of tools to resolve rRNA processing and degradation kinetics across the entire maturation pathway. Traditional methods, including pulse-chase radiolabeling^25,26^ and northern blotting,^27^ were instrumental in defining the pre-rRNA processing pathway by tracking the temporal abundance of intermediates.^1^ Although they sensitively monitor intermediate dynamics within an experiment, comparisons across cellular conditions are difficult because labeling efficiency, RNA recovery, and loading normalization vary between samples. Moreover, it has been difficult to determine whether reduced precursor abundance reflects cleavage into downstream intermediates or increased degradation, as both processes decrease the abundance of a given species. Resolving this distinction is essential for understanding how rRNA maturation is altered under oncogenic drive. Consequently, despite many studies reporting increased rRNA transcription and elevated ribosome abundance in cancer cells,^4,5,24,28^ it remains unknown whether ribosome production is coordinated across maturation steps or whether specific stages act as bottlenecks that limit ribosome output. Addressing this gap requires quantitative, pathway-wide measurements that resolve transcription, processing, and degradation kinetics, to determine how the elevated output of functional ribosomes is achieved and potentially constrained under oncogenic conditions.

## Results

### MYC activation reduces ribosome biogenesis efficiency

To determine whether oncogene-driven increases in pre-rRNA transcription produce a proportional increase in ribosomes (**Fig. 1a**), we first quantified the ribosome biogenesis yield (efficiency) by measuring pre-rRNA transcription and newly assembled cytoplasmic ribosomes. We quantified transcription rate by labeling nascent RNAs with a uridine analog, 5-ethynyl uridine (5eU). After a 30-minute pulse, newly transcribed RNA was visualized by conjugating 5eU to a fluorophore via click chemistry (**Fig. 1b**).^29–32^ The nucleolar 5eU signal was used as a readout of rRNA transcription, the predominant transcriptional activity in the nucleolus.^32,33^

**Fig. 1:**
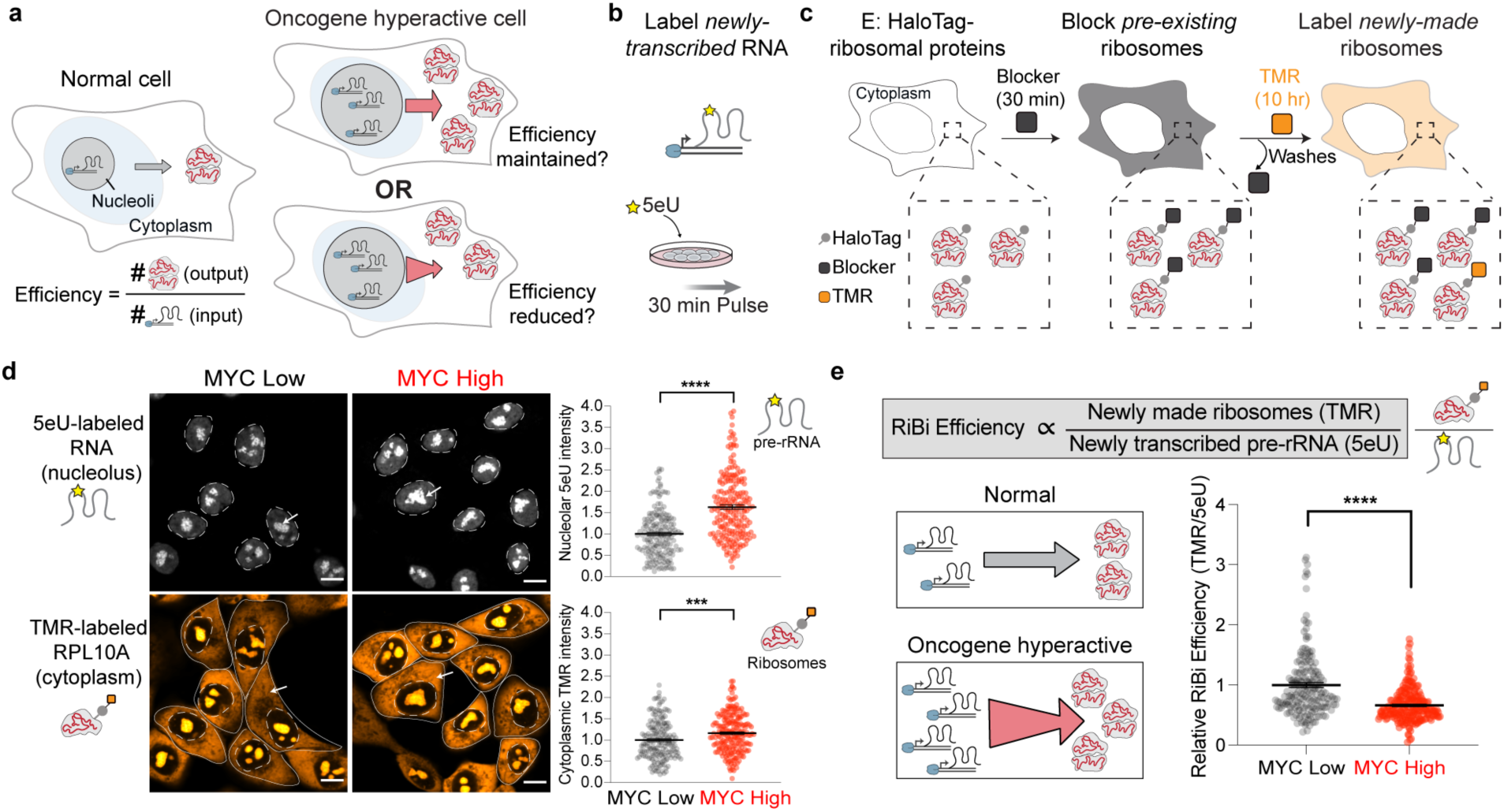
Efficiency of ribosome biogenesis is reduced upon oncogene activation. **a**, Conceptual framework for ribosome biogenesis efficiency. In normal cells (left), precursor rRNAs (pre-rRNAs) are transcribed in the nucleolus and assembled ribosomes are exported to the cytoplasm. Ribosome biogenesis efficiency is defined as the ratio of ribosome output to pre-rRNA transcription input. Upon oncogene activation (right), increased pre-rRNA transcription may either be accommodated by downstream processing steps, maintaining proportional ribosome production (top), or exceed processing capacity, resulting in reduced ribosome output relative to input and therefore decreased efficiency (bottom). **b**, Cells are pulsed with 5eU (30 min) to label newly-transcribed RNA for quantifying transcription rate. **c**, Cells with endogenously tagged (E:) HaloTag-ribosomal proteins (RPs) are labeled with a non-fluorescent HaloTag ligand “Blocker” (30 min) to block all the pre-existing ribosomes. After multiple washes, cells are then incubated with a fluorescent HaloTag ligand “TMR” (10 h) to label the newly-made ribosomes. **d**, Example images of MCF10A cells (E: HaloTag-RPL10A) with both 5eU-labeled rRNA (white) and TMR-labeled (orange) RPL10A in MYC-low and MYC-high conditions. Both signals were measured in the same cells, enabling per-cell quantification (in e). Top right, quantified 5eU intensity in the nucleolus for rRNA transcription rate. Bottom right, quantified TMR intensity in the cytoplasm for nascent ribosome production rate. *** p value = 0.0003; **** p value <0.0001 (two-tailed Mann-Whitney test); n = 211 (MYC-high), 181 (MYC-low) cells. Scale bars = 10 μm. **e**, Ribosome biogenesis (RiBi) efficiency is quantified as the ratio of cytoplasmic TMR (HaloTag-RPL10A) to pre-rRNA transcription (measured by 5eU) per cell. Left, schematic illustrating proportional coupling between transcription and ribosome output in normal cells and inefficient propagation in oncogene-hyperactive cells. Right, distribution of relative RiBi efficiency (TMR/5eU) in MYC-low and MYC-high cells. ****p < 0.0001 (two-tailed Mann-Whitney test); n = 211 (MYC-high), 181 (MYC-low) cells. Black lines denote mean values; error bars are s.e.m, in both d and e.

To quantify ribosome biogenesis output, we endogenously tagged ribosomal proteins RPL10A and RPS14 with HaloTag (**Extended Data Fig. 1a**), enabling covalent labeling of newly assembled ribosomes.^34,35^ Briefly, a non-fluorescent blocker ligand was first added to covalently block all pre-existing ribosomes, followed by addition of a fluorescent ligand (Tetramethylrhodamine, TMR) to selectively mark nascent ribosomes during a defined labeling period (**Fig. 1c**). During this time, newly synthesized RPL10A or RPS14 proteins were fluorescently labeled with TMR, incorporated into assembling pre-ribosomal particles in the nucleolus, and subsequently exported to the cytoplasm for maturation. Polysome profiling showed that cytoplasmic TMR signal predominantly co-sediments with assembled ribosomal subunits or ribosomes rather than free ribosomal proteins (**Extended Data Fig.1b**). We therefore measured cytoplasmic TMR signal as a proxy for newly assembled ribosomal particles in the cytoplasm (**Fig. 1d**).

Measurements of rRNA transcription (5eU) and ribosome production (TMR) can be obtained by imaging the same cells, enabling quantification of the relationship between rRNA transcription and ribosome output, at single-cell resolution. Using this approach, we quantified ribosome production efficiency in immortalized human breast epithelial MCF10A^36,37^ cells harboring an inducible MYC overexpression system (**Extended Data Fig. 1c**). In this Tet-Off system, doxycycline (Dox) keeps the exogenous MYC in an off state (hereafter referred to as “MYC-low”), while Dox withdrawal induces MYC overexpression (hereafter “MYC-high”). Unless otherwise indicated, cells were maintained in the MYC-high state for 72 h prior to measurement. We confirmed that MYC overexpression does not alter 5eU incorporation efficiency, so 5eU signal directly reflects transcription rate (**Extended Data Fig. 1d**). Consistent with MYC’s established role in ribosome biogenesis,^24,38^ MYC overexpression increased both nucleolar pre-rRNA transcription and cytoplasmic ribosome production (**Fig. 1d**). Strikingly, however, the increase in cytoplasmic ribosome production was substantially less than the increase in pre-rRNA transcription, revealing a pronounced reduction in ribosome biogenesis yield upon MYC induction. This reduction was observed for both SSU (RPS14) and LSU (RPL10A) by single-cell volumetric imaging analysis (**Fig. 1d-e and Extended Data Fig. 1e-f**) and by bulk biochemical quantification of 5eU and TMR signals (**Extended Data Fig. 1g-l**).

### Kinetic mapping reveals delayed late-stage rRNA processing

The observed reduction in overall ribosome biogenesis yield led us to hypothesize that this inefficiency arises from alterations in specific rRNA processing steps. To test this, we applied 5eU-sequencing (5eU-seq), a pulse-chase RNA labeling and purification method^39–41^ that we recently optimized to selectively isolate newly synthesized pre-rRNAs while minimizing background from abundant mature rRNAs.^31^ Briefly, newly transcribed RNAs were pulse-labeled with 5eU and chased with excess uridine, allowing time-resolved tracking of rRNA maturation. Sequencing of labeled RNA resolved the temporal dynamics of pre-rRNA cleavage and modification (2’-*O*-methylation) (**Fig. 2a**). Cleavage sites were classified as early, middle, or late according to their timing^31^ and established order in the rRNA maturation pathway^42^ (**Fig. 2a**). This approach allowed us to quantify the fraction of transcripts cleaved at each site (**Fig. 2b)**. We note that productive cleavage occurs only after successful rRNA folding, ribosomal protein incorporation, and remodeling during ribosome assembly, such that the timing of cleavage also reflects the progression of upstream assembly steps.^2,11^

**Fig. 2:**
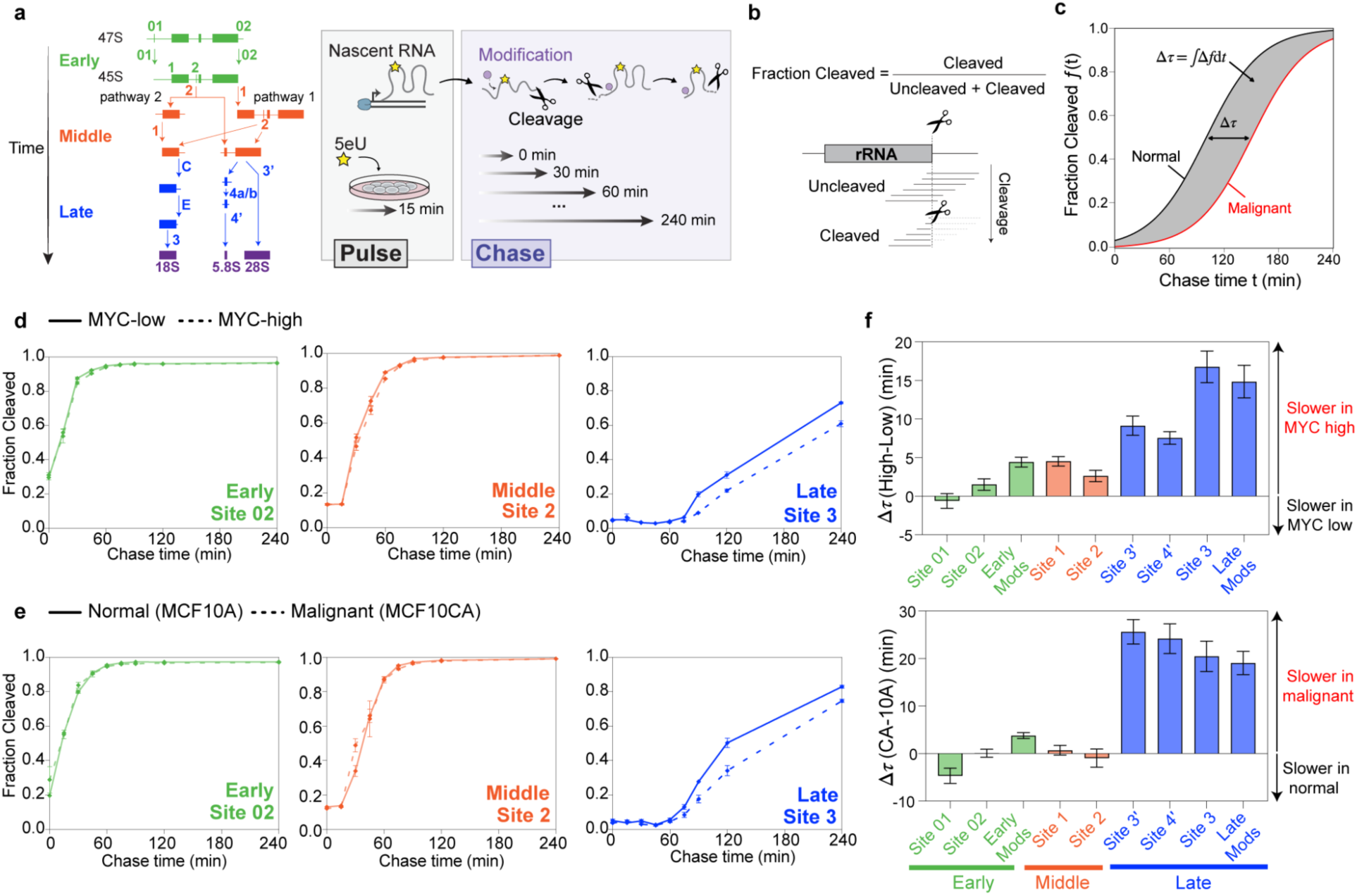
Kinetic mapping reveals delayed late-stage rRNA processing. **a**, Left, schematic of pre-rRNA cleavage steps classified as early (green), middle (red) and late (blue) based on their order in the cleavage pathway,^6,42^ adapted from ref. 31. Right, 5eU pulse chase followed by sequencing (5eU-seq). Cells are pulsed with 5eU for 15 min to label newly transcribed RNA and then chased over 0-240 min to measure cleavage and modification of pre-rRNA over time via sequencing. **b**, The “fraction cleaved” metric is calculated by quantifying the number of “cleaved” reads (ending at a cut site) divided by the total number of reads (uncleaved (spanning a cut site) + cleaved). **c**, Schematic of delay time Δτ calculation, which is computed by the time integral of the difference in fraction cleaved between conditions (shaded area). See Methods for details. **d-e**, Fraction of pre-rRNA cleaved at early, middle, and late sites over time. n = 2 per timepoint. d, Solid lines are MYC-low cells and dashed lines are MYC-high cells. e, Solid lines are normal (MCF10A) cells and dashed lines are malignant (MCF10CA) cells. **f**, Quantified delay time for all cleavage and modification (Mods) sites between MYC-high and MYC-low (top panel) or MCF10CA and MCF10A (bottom panel). See raw data in Extended Data Fig. 2. Error bars indicate propagated uncertainty of the Δτ estimate, see Methods.

Using this approach, we systematically quantified the kinetics of pre-rRNA processing upon oncogene activation. In addition to the inducible MYC system, we also compared rRNA processing dynamics between the normal-like breast epithelial cell line MCF10A and its malignant derivative MCF10CA1a.cl1 (hereafter referred to as “MCF10CA”), generated by HRAS^G12V transformation and subsequent *in vivo* selection.^43,44^ 5eU-seq revealed that while early (site 01, site 02) and middle (site 1 and site 2) cleavage steps proceed with comparable kinetics in normal, MYC-high and malignant cells, late (site 3’, site 4’, and site 3) cleavage steps are markedly delayed in MYC-high (**Fig. 2d and Extended Data Fig. 2a**) and malignant (MCF10CA) cells (**Fig. 2e and Extended Data Fig. 2b**). The slowing down of these cleavage steps can be quantified by a delay time Δτ, which is the difference in average cleavage time between conditions for each processing site (**Fig. 2c and Methods**). Using this metric, we found that late cleavage steps are consistently slower in both MYC-overexpressing and malignant (MCF10CA) cells compared to normal (MCF10A) cells (**Fig. 2f**). Consistent with these cleavage delays, 2’-*O*-methylation events during late pre-rRNA processing, such as 28S-Um4498 and 28S-Gm4499, were also slower in MYC-high and malignant cells (**Fig. 2f and Extended Data. Fig. 2c**). Taken together, kinetic mapping using 5eU-seq reveals a selective delay in late-stage pre-rRNA processing under oncogenic conditions.

### Quantifying pre-rRNA degradation with spike-in normalized 5eU-seq

Having established that late rRNA processing steps are delayed in cancer, we next asked whether these delays are accompanied by increased degradation of accumulating pre-rRNA intermediates. Testing this hypothesis required a quantitative measurement of pre-rRNA degradation by tracking its abundance over time. We therefore incorporated external RNA spike-in standards into the 5eU-seq workflow, adapting principles from established metabolic RNA turnover approaches^40,45,46^ (**Fig. 3a**, **Extended Data Fig. 3a and Methods**). We validated this normalization strategy using mock 5eU-labeled samples, confirming accurate spike-in-based quantification (**Extended Data Fig. 3b-d**). Moreover, spike-in normalization reliably captured the degradation kinetics of known short-lived and long-lived RNAs^46–48^ (**Extended Data Fig. 3e and Supplementary Note 2**). Using this approach, we observed a greater reduction in the abundance of 28S- and 18S-containing pre-rRNAs during chase in malignant (MCF10CA) cells compared to normal cells (**Fig. 3b**), indicating increased degradation of pre-rRNA intermediates. These data suggest that part of the incoming flux of rRNA intermediates is degraded, consistent with delayed processing representing kinetic bottlenecks that reduce the overall yield of ribosome production.

**Fig. 3:**
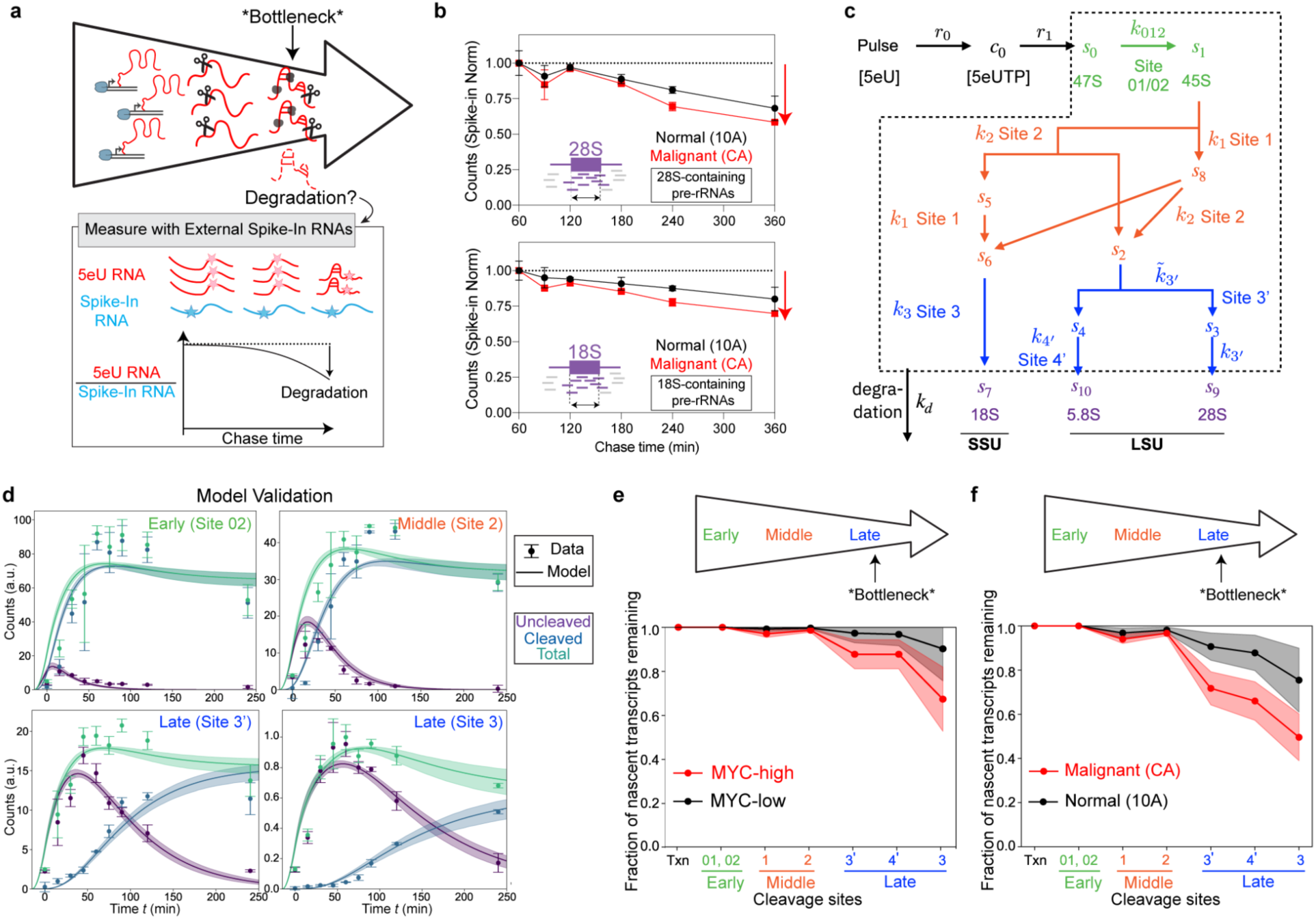
Late-stage pre-rRNA loss creates bottlenecks in ribosome biogenesis. **a**, Spike-in–normalized quantification of pre-rRNA degradation. External spike-in RNAs were added in proportion to total RNA for each sample during the 5eU-seq workflow (Methods and Extended Data Fig. 3a), enabling normalization of 5eU-labeled pre-rRNA abundance across chase timepoints to quantify loss by degradation. **b**, Decay of 28S-containing (top) and 18S-containing (bottom) pre-rRNA intermediates over time measured by region count analysis (see Methods), for MCF10A (black) and MCF10CA (red) cells. Error bars are s.e.m. **c**, A minimal mathematical model for pre-rRNA maturation, which describes 5eU conversion to 5eUTP, rRNA transcription, sequential cleavage at defined processing sites, and degradation of intermediates (each with rate k_d_) across the ribosome biogenesis assembly line. **d**, Model validation in MCF10CA cells (with data not used for fitting): Model-predicted dynamics (lines) of cleaved, uncleaved and total (cleaved+uncleaved) species at each cleavage step compared with independent experimental spike-in-normalized measurements (points, mean ± s.d., n=2 per time point). See Extended Data Fig. 4 for model validation in MCF10A, and Supplementary Note 2 for validation in MYC-low and MYC-high cells. **e-f**, Model-derived fraction of nascent transcripts remaining along the rRNA processing pathway in MYC-low vs MYC-high (e) or MCF10A vs MCF10CA (f). Y axis, the fraction of newly transcribed pre-rRNA molecules that have undergone the corresponding cleavage without prior degradation (normalized to transcription). In d–f, Shaded regions indicate uncertainty from model fitting (see Supplementary Note 2). MYC high and malignant cells show a pronounced reduction in efficiency at late processing steps, corresponding to the bottlenecks indicated in the schematic above.

### Mathematical modeling deconvolves cleavage and degradation to quantify stepwise processing efficiency

So far, we have shown that oncogene activation leads to elevated rRNA transcription, slower late-stage pre-rRNA processing, and increased degradation of pre-rRNA intermediates. However, rRNA maturation is an interconnected pathway in which measurements at any cleavage site reflect cumulative upstream processing.^2,11,49^ In addition, transcription, cleavage, and degradation occur concurrently during 5eU pulse-chase labeling. As a result, disentangling how and to what degree specific rRNA processing and degradation steps are modulated in hyperproliferative cells is challenging. We thus sought to gain a deeper understanding of the process by analyzing our data in the framework of a mathematical model, which can also take into account potentially confounding effects of non-instantaneous 5eU incorporation and washout (**Supplementary Note 1**).

We considered a minimal chemical kinetic model that describes the dynamics of 5eU-labeled pre-rRNA, including intracellular 5eU to 5eUTP conversion, transcription, cleavage, and degradation (**Fig. 3c and Supplementary Note 2**). During the pulse, cells continuously uptake 5eU and convert it to 5eUTP, which is incorporated into nascent pre-rRNA by transcription both during and after the pulse. The model assumes nascent transcripts are cleaved with linear kinetics at all sites, and intermediates are susceptible to degradation prior to completion of all cleavage steps, whereas the fully processed product is protected from degradation. These processes are described by a system of ordinary differential equations, whose model parameters are inferred by maximizing the likelihood of the observed data. To assess whether the minimal model is able to capture the measured pre-rRNA dynamics, we fit the model to a subset of the data and evaluated its ability to predict the remaining measurements (**Supplementary Note 2**). Specifically, by fitting the model to the time-course measurements of the fraction of transcripts cleaved at each site, as well as the decay of 28S- or 18S-containing pre-rRNAs, we inferred the cleavage and degradation rates, in MCF10A/MCF10CA cells (**Extended Data Fig. 4a-c**) and the inducible MYC overexpression system (**Extended Data Fig. 4f-h**). The best-fit parameters revealed that malignant (or MYC-high) cells have higher transcription activity (consistent with **Fig. 1d**), lower late-stage processing rates, and higher degradation rates than normal (or MYC-low) cells.

Next, we evaluated the model’s predictive performance by using the inferred parameters to predict the dynamics of both cleaved and uncleaved pre-rRNAs at each cleavage site, which were not used for fitting. The predictions were in agreement with the measured spike-in normalized cleaved and uncleaved counts (**Fig. 3d, Extended Data Fig. 4e and Supplementary Note 2**), demonstrating that our minimal model provides a good description of pre-rRNA processing dynamics.

### Late-stage pre-rRNA loss creates bottlenecks in ribosome biogenesis

By accounting for both cleavage and degradation, our mathematical model quantifies, at each cleavage step, the fraction of the original transcripts that reach that step and undergo cleavage. In both MYC-low and MYC-high cells, most transcripts completed early and middle-stage processing with minimal degradation, whereas the majority of degradation occurred during late-stage processing, and was more pronounced in MYC-high cells. Accordingly, the fraction of nascent pre-rRNA completing the final cytoplasmic cleavage step (site 3) decreased from 90 ± 14% (mean ± s.d.) in MYC-low cells to 67 ± 14% in MYC-high cells (**Fig. 3e and Extended Data Fig. 4i**). To compare the model-derived estimate with experimental measurements, we calculated a relative ribosome biogenesis efficiency by normalizing the fraction of transcripts completing site 3 processing in MYC-high cells to MYC-low cells (**Supplementary Note 3**). This yielded an estimated 25% ± 16% reduction in ribosome biogenesis efficiency upon MYC activation (**Fig. 3e**). Consistent with this estimate, independent measurements using 5eU labeling of nascent rRNA transcription and TMR labeling of newly synthesized ribosomal subunits revealed a similar decrease in efficiency (29 ± 10% for LSU via HaloTag-RPL10A and 25 ± 10% for SSU via HaloTag-RPS14) (**Extended Data Fig. 1e-l**). Applying the same analysis to the MCF10A/MCF10CA system revealed a comparable shift toward increased late-stage degradation, with the fraction of transcripts reaching site 3 decreasing from 80 ± 15% in normal MCF10A cells to 52 ± 11% in malignant MCF10CA cells, corresponding to an estimated 35% ± 10% reduction in ribosome biogenesis efficiency (**Fig. 3f, Extended Data Fig. 4d and Supplementary Note 4**). These results demonstrate that delayed late-stage processing is tightly coupled to increased degradation of pre-rRNA intermediates, further suggesting that imperfectly upregulated processing gives rise to kinetic bottlenecks that limits ribosome biogenesis efficiency in oncogene-active cells.

### Late-stage perturbations selectively reduce ribosome biogenesis efficiency in MYC-high cells

Our data support a model in which late-stage processing impedes efficient ribosome production during oncogene activation, such that perturbations at these steps would have a disproportionately large impact on ribosome biogenesis in cancer cells. Indeed, our theoretical model predicts that decreasing the cleavage rate at late site 3’ or site 3 leads to a more substantial reduction in yield in MYC-high or malignant cells compared to MYC-low or normal cells (**Supplementary Note 2**). Because ribosome biogenesis factors are essential and complete loss would impair viability across all conditions, we performed partial knockdown to reduce their levels without complete depletion. We then tested this prediction experimentally by measuring ribosome biogenesis efficiency using the 5eU and TMR approach described in Fig. 1, following knockdown of early, middle, or late ribosome biogenesis factors in the inducible MYC system (**Supplementary Note 5**). Knockdown of late processing factors, including PES1, which is required for ITS2 processing during LSU maturation, and NOB1, the cytoplasmic endonuclease that performs the final cleavage at site 3 to generate mature 18S rRNA,^11,50^ caused a significantly greater reduction in efficiency in MYC-high cells, consistent with model predictions. In contrast, depletion of early (KRI1; co-transcriptional processing) or middle (UTP20; 5′ ETS processing) factors^50^ produced comparable reductions in both MYC-high and MYC-low conditions (**Extended Data Fig. 5a-e**).

### Targeting ribosome biogenesis bottlenecks suppresses MYC-driven cell and tumor growth

Given the selective impairment of ribosome biogenesis efficiency in MYC-high cells upon bottleneck perturbation, we hypothesized that this vulnerability could be selectively targeted. To test this, we monitored the proliferation of MYC-low and MYC-high cells in 2D cell culture following perturbations of ribosome biogenesis (**Fig. 4a-b**). Knockdown of early or middle ribosome biogenesis factors had comparable effects on MYC-low vs MYC-high cells, whereas perturbation of late-stage factors preferentially suppressed the proliferation of MYC-high cells (**Fig. 4c and Extended Data Fig. 6a-b**).

**Fig. 4:**
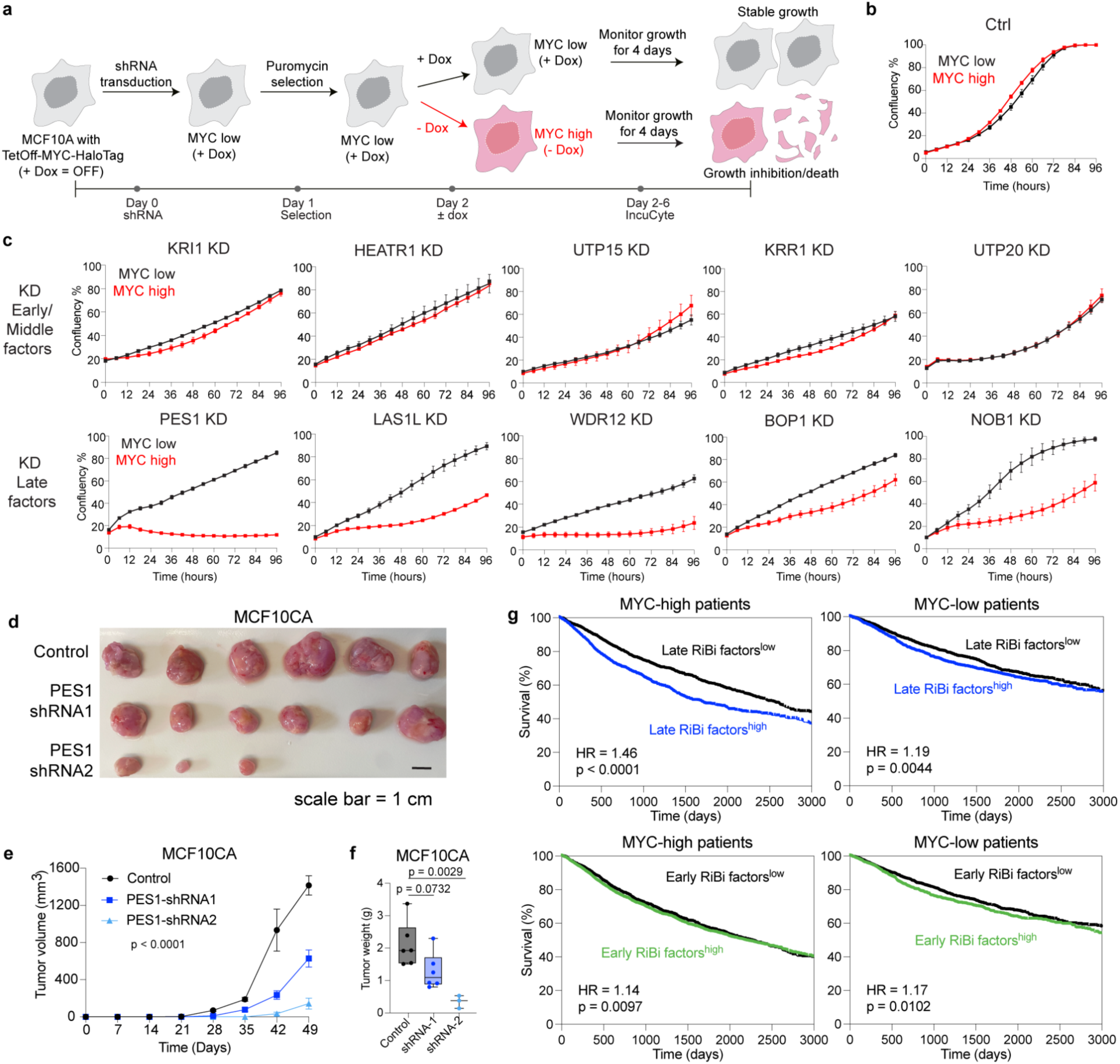
Targeting ribosome biogenesis bottlenecks suppresses MYC-driven cell and tumor growth. **a**, Experimental workflow to test MYC-dependent sensitivity to ribosome biogenesis perturbation. MCF10A cells with TetOff MYC-HaloTag were transduced with shRNAs targeting ribosome biogenesis (RiBi) factors and selected with puromycin. MYC expression was controlled by doxycycline (Dox): MYC-low (+ Dox) and MYC-high (- Dox). Cell proliferation was monitored for 4 days by live-cell imaging (IncuCyte). **b**, Growth of MYC-high and MYC-low cells with control shRNAs. **c**, Effects of RiBi factor knockdown on cell proliferation. Time-resolved confluency measurements following knockdown (KD) of early/middle rRNA processing factors (KRI1, HEATR1, UTP15, KRR1, UTP20) or late factors (PES1, LAS1L, WDR12, BOP1, NOB1). Black, MYC-low cells; red, MYC-high cells. Points represent mean ± s.e.m. n=3 per time point. **d-f**, Growth of MCF10CA xenograft tumors following PES1 knockdown. d, Representative images of endpoint tumors excised from female NSG mice injected with control cells or cells expressing PES1 shRNAs (shRNA1 or shRNA2). Scale bar = 1 cm. e, Tumor volume over time. Error bars represent mean ± s.e.m. Tumor growth curves were compared using two-way repeated-measures (RM) ANOVA with Geisser-Greenhouse correction; P values indicate the time x treatment interaction. f, Tumor weight at endpoint. Box plots show medians (lines), boxes (25th-75th percentiles), and whiskers (min-max). Statistical significance was assessed using one-way ANOVA with Tukey’s test for multiple comparisons. n = 7 (MCF10CA control); 6 (MCF10CA PES1-shRNA1); 3 (MCF10CA PES1-shRNA2) mice for tumor growth curves. One mouse from the control group died before endpoint tumor collection and was therefore excluded from tumor weight measurements. Final tumor weight analyses were performed with n = 6, 6 and 3 mice. **g**, Clinical relevance of ribosome biogenesis bottlenecks in patients. Kaplan–Meier analysis of overall survival among TCGA patients (n = 9,642). Patients were first stratified by MYC expression (high vs low, median split) and subsequently by early or late ribosome biogenesis (RiBi) factor expression (high vs low, median split) based on mRNA expression measured in primary tumors. Hazard ratios (HR) were estimated using a multivariable Cox proportional hazards regression model including MYC status, RiBi factor expression, and a MYC×RiBi interaction term. Reported HRs represent MYC-adjusted effects of RiBi factor expression on survival within each MYC stratum. P values were calculated using two-sided Wald tests.

We next asked whether the susceptibility of late-stage ribosome biogenesis bottlenecks extends to tumor models. MYC pathway activation is estimated to occur in up to 70% of tumors,^17,51,52^ particularly elevated in basal-like and triple-negative breast cancers, aggressive cancer subtypes associated with poor clinical outcomes and few targeted therapeutic options.^53–55^ We thus examined the triple-negative breast cancer line SUM159,^56,57^ and also MCF10CA cells, both of which displayed elevated MYC level relative to MCF10A (**Extended Data Fig. 7a**). Consistent with the sensitivity observed in cell culture, knockdown of a representative late-stage ribosome biogenesis factor, PES1, using two independent shRNAs significantly reduced tumor growth in both MCF10CA and SUM159 xenografts in female NSG mice (**Fig. 4d-f and Extended Data Fig. 7b-e**). PES1 has also been identified in prior functional genomic screens as a candidate synthetic lethal target in MYC-overexpressing cells;^58^ here, we show that this vulnerability arises from late-stage ribosome biogenesis bottlenecks.

Analysis of TCGA pan-cancer patient cohorts further supported the clinical relevance of late-stage ribosome biogenesis bottlenecks. High expression of late ribosome biogenesis factors was associated with reduced patient survival, particularly in MYC-high tumors, whereas early factors showed weaker associations (**Fig. 4g; see Supplementary Table 1 for the list of early and late factors**). Collectively, these findings identify late-stage ribosome biogenesis as a selective vulnerability in MYC-driven tumors.

## Discussion

Ribosome biogenesis is broadly upregulated in cancer, yet whether this increase is proportionally coordinated across rRNA transcription, processing, and assembly has remained unclear. By developing a quantitative framework that combines pulse-chase mapping with mathematical modeling, we resolved pre-rRNA transcription, processing, and turnover dynamics to measure ribosome biogenesis efficiency and systematically identify rate-limiting steps along the course of rRNA maturation. We find that oncogene activation creates a bottleneck in late-stage processing, leading to delayed maturation and increased degradation of pre-rRNA intermediates (**Fig. 5a**). Consistent with this, reducing late ribosome biogenesis factors suppresses MYC-driven cell proliferation and tumor growth (**Fig. 5b**). Oncogene activation therefore creates bottlenecks within the ribosome assembly pathway, exposing vulnerabilities that could be therapeutically exploited. This inefficiency challenges the prevailing assumption that enlarged nucleoli and high rRNA transcription directly reflect productive ribosome output, and instead suggests that cancer cells sustain growth despite substantial losses along the ribosome assembly pathway.

**Fig. 5:**
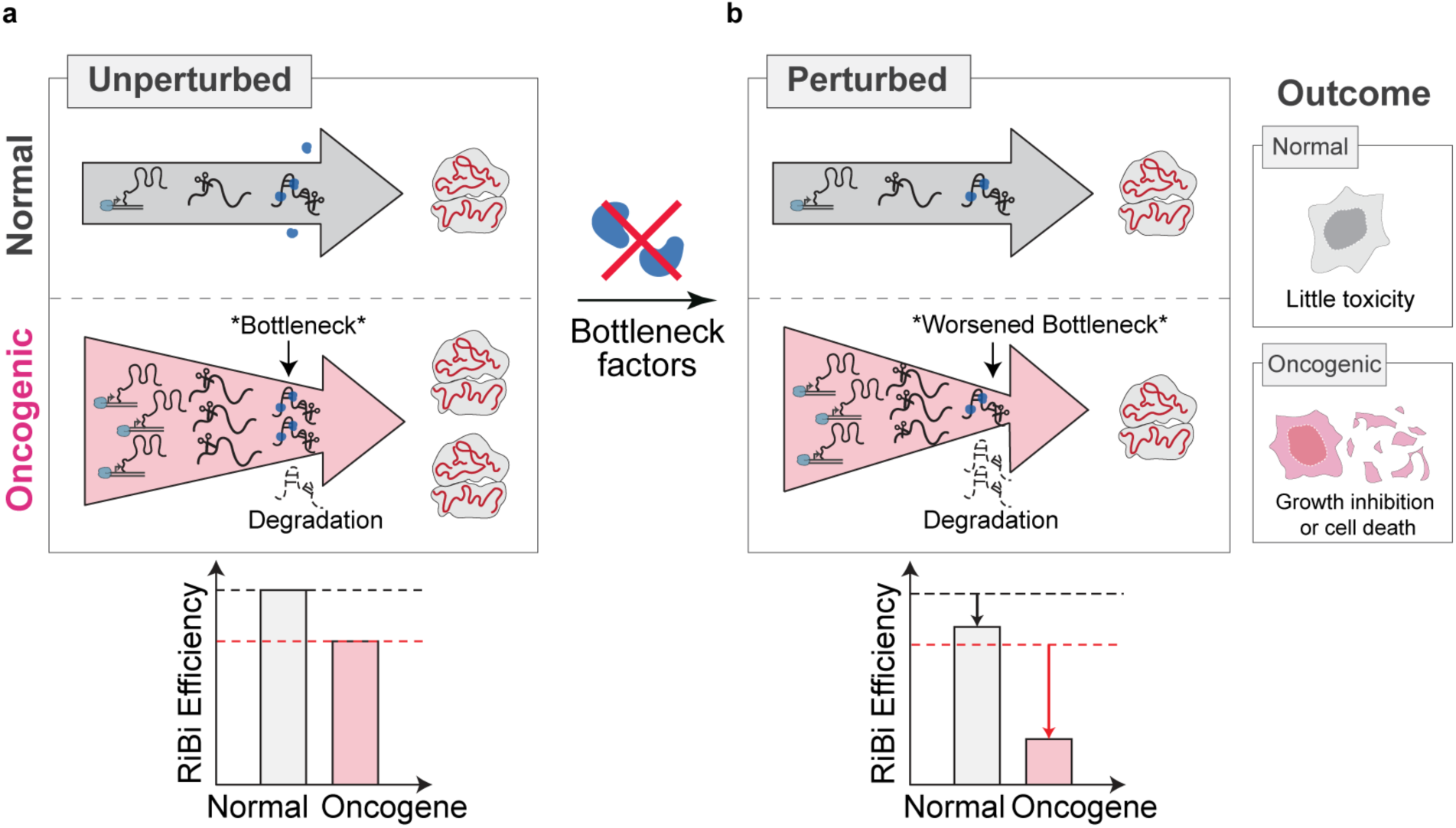
Oncogene activation creates a bottleneck in ribosome biogenesis that confers selective vulnerability. **a**, Conceptual model. In normal cells, rRNA transcription, processing, and ribosome assembly proceed in a balanced manner, allowing efficient ribosome production. Upon oncogene activation (e.g., MYC), transcription increases markedly, but downstream processing capacity does not scale proportionally, creating a late-stage ribosome biogenesis (RiBi) bottleneck. As a result, excess pre-rRNA accumulates and a fraction is degraded, lowering overall ribosome biogenesis efficiency despite elevated transcription. **b**, Partial inhibition of bottleneck steps has minimal effects in normal cells but worsens the bottleneck in oncogene-active cells, increasing pre-rRNA turnover, reducing productive ribosome output, and selectively impairing growth or viability.

A central implication of our findings is that ribosome biogenesis in cancer is not simply elevated, but fundamentally imbalanced. Rather than proportionally increasing productive output, heightened ribosome biogenesis reflects a trade-off: increased biosynthetic flux is achieved at the cost of reduced yield, placing tumor cells closer to the limits of processing and quality-control capacity. Sustained rRNA overproduction, coupled with limited downstream processing capacity, likely increases the burden on surveillance and degradation pathways that remove defective or stalled assembly intermediates. The molecular machinery responsible for recognizing, processing, and recycling these aberrant pre-rRNAs remains incompletely defined.^59–62^ Systematic identification of such quality-control factors, and assessment of whether cancer cells exhibit increased dependence on these pathways, will be an important direction for future work.

Efforts to therapeutically target ribosome biogenesis have been constrained by poor selectivity and substantial toxicity, largely because existing strategies globally suppress rDNA transcription or nucleolar function.^63,64^ A key challenge is that many ribosome biogenesis factors are essential for normal cell viability, such that complete inhibition is broadly deleterious. Our findings instead suggest that selective vulnerability may arise from partial perturbation of rate-limiting processing steps in oncogene-driven contexts, where cells operate closer to the limits of processing capacity. In this regime, modest reductions in processing activity can disproportionately impair ribosome production in oncogene-driven cells. Achieving this level of selective tuning may nevertheless be challenging, as the therapeutic window for partially inhibiting essential factors is likely to be narrow and context-dependent.

More broadly, these findings support a general principle in cancer biology: oncogenic hyperactivation of biosynthetic pathways can create intrinsic capacity limits that are not apparent under normal growth conditions, thereby generating selective liabilities.^65–67^ Targeting such bottlenecks, rather than upstream drivers alone, may therefore provide a complementary strategy for exploiting the systems-level stresses imposed by oncogenic growth programs.

The framework developed here provides a quantitative approach for dissecting coordination across multistep biosynthetic pathways. Applying similar kinetic analyses across different oncogenic drivers, tumor types, or physiological contexts may reveal whether distinct cell growth programs generate unique bottlenecks in ribosome biogenesis. Together, our findings highlight that highly coordinated cellular processes may operate near intrinsic capacity limits. Identifying such capacity limits may therefore provide a general strategy for revealing vulnerabilities created by imbalances in complex biosynthetic pathways across physiology and disease.

## Methods

### Statistics and Reproducibility

Statistical analyses were performed using GraphPad Prism 10 unless otherwise stated. Statistical tests, p-values, and sample sizes are specified in figures or figure legends. P value less than 0.05 was considered statistically significant. Microscopy imaging, western blotting and RT-qPCR were repeated independently at least three times and/or using at least three biologically independent samples.

### Cell culture

Cells were cultured at 37 °C with 5% CO_2_ in a humidified chamber. All cell culture media were supplied with 1% streptomycin and penicillin (GIBCO, 15140122). MCF10A (RRID: CVCL_0598) and MCF10CA1a.cl1 (CVCL_6676) cells were cultured in DMEM/F12 medium (Thermo Scientific, 11320082) supplied with 5% horse serum (Sigma, H1138), 20 ng/mL EGF (Novoprotein, C029), 10 ng/mL insulin (Sigma, 91077C), and 1 µg/mL hydrocortisone (Sigma, H0888). HEK293T (CVCL_0063, ATCC) cells were cultured in DMEM (GIBCO, 11995065) supplied with 10% FBS (R&D Systems, S11150H). SUM159 (RRID: CVCL_5423) was cultured with Ham’s F12 (Corning, 10-080-CV) supplemented with 10% FBS (R&D Systems, S11150H), 1 µg/mL hydrocortisone (Sigma, H0888) and 10 ng/mL insulin (Sigma, 91077C). All cells were routinely checked for mycoplasma contamination and authenticated.

### Microscopy

Images were acquired on a Nikon CSU-W1 SoRa spinning-disk confocal microscope (Yokogawa) capable of pixel-reassignment super-resolution imaging. A Nikon CFI Plan Apo Lambda 60x oil objective (NA 1.40; MRD71670) was used for all experiments. Depending on the experiment, images were collected either in conventional CSU-W1 confocal mode or in SoRa super-resolution mode (2.8x magnification). Excitation was provided by 405, 488, 561 and 640 nm laser lines. Z-stacks were acquired using a Mad City Labs piezo Z stage with 0.7 µm step size over a total depth of 20 µm.

### Western blotting

Western blotting was performed following standard procedures. Cells were harvested from 6-well plates by trypsinization and washed twice in PBS, followed by lysis in 80-120 µL RIPA buffer (ThermoFisher, 89900) supplemented with protease and phosphatase inhibitors (1:100, Thermo Scientific, PI78440) and benzonase (1:300, Sigma Aldrich, E8263-25KU). Lysates were incubated on ice for 15–20 min and clarified by centrifugation at 10,000g for 10 min at 4°C. Protein concentration was quantified using the Pierce Bradford assay (ThermoFisher, 23246). Equal amounts of protein (13-15 µL) were mixed with NuPAGE LDS sample buffer (ThermoFisher, NP0004) and sample reducing agent (ThermoFisher, NP0007), denatured at 95°C for 5 min, and resolved on NuPage 4–20% Bis-Tris gels (ThermoFisher, NP0322BOX) using MES running buffer (ThermoFisher, NP0002). Proteins were transferred to nitrocellulose or PVDF membranes using the Trans-Blot Turbo Transfer System (BioRad, 1704150) and blocked in 5% BSA in TBST. Membranes were incubated with primary antibodies (**Supplementary Table 1**) overnight at 4°C, washed, and incubated with HRP-conjugated secondary antibodies (**Supplementary Table 1**) for 45 min at room temperature. Blots were developed using the clarity western ECL substrate (BioRad, 1705061) and imaged within 10 min of reagent application.

### RNA isolation

For each well of a 6-well plate, cells were lysed in 200-300 μL of 1× Buffer RLT (QIAGEN, 79216), and total RNA was purified using the QIAGEN RNeasy Mini Kit (74104). Residual genomic DNA was removed by treating the isolated RNA with TURBO DNase (Thermo Fisher Scientific, AM2238) for 30 min to 1 h at 37 °C. Following DNase digestion, RNA was further purified and concentrated using the Zymo RNA Clean and Concentrator-25 kit (R1017).

### Endogenous tagging of ribosomal proteins

N-terminal endogenous tagging of ribosomal proteins (RPS14 and RPL10A) was conducted as previously described.^31,68,69^ An oligonucleotide pair encoding an RPS14-targeting (CTCAGAAATGGCACCTCGAA) or RPL10A-targeting (TTCAGGACCAACTCACCTCA) gRNA was cloned into a modified lentiCRISPRv2-puro plasmid (courtesy of Aaron Lin) via the BsmBI restriction site. One plasmid containing gRNA and one plasmid with donor sequence for homology-directed repair (HDR) were transfected into the MCF10A cells using the FuGENE® HD Transfection Reagent following the manufacturer’s instructions (Promega, E2311). The donor plasmids were constructed by cloning the HaloTag with a flexible linker flanked by 300-500 bp homology arms complementary to the N-terminus of the RPS14 or RPL10A coding sequence into the pUC19 vector (ThermoFisher, SD0061) (**Supplementary Table 1**). Three days after transfection, cells were labeled with HaloTag-JF646 ligand (Janelia) to sort for HaloTag-positive single cells into 96-well plates. After expansion of these single-cell clones, clones were screened and validated through western blotting and junction PCR (primers in **Supplementary Table 1**) of the specific genomic locus (**Extended Data Fig. 1a**).

### Reverse transcription quantitative PCR

Reverse transcription quantitative PCR (RT-qPCR) was performed on an Applied Biosystems QuantStudio 3 Real-Time PCR System instrument (Thermofisher, A28567). All primers were synthesized from IDT and primer sequences are listed (**Supplementary Table 1**). RT-qPCR reaction mixtures were prepared using the Luna Universal One-Step RT-qPCR Kit (NEB, E3005) following manufacturer’s instructions.

### Lentiviral packaging

HEK293T cells were seeded one day before transfection in a 10 cm plate to reach 70-80% the day of transfection. The desired plasmids were transfected together with helper plasmids VSVG and PSP via lipofectamine^TM^ 3000 (Invitrogen, L3000008) following standard procedures. Briefly, 75 µl Opti-MEM™(Thermofisher, 31985062), 2400 ng of VSVG, 7200 ng of PSP, 2400 ng of desired plasmids and 45 µl of p3000 were added in order into one appendorf tube, while 1500 µl Opti-MEM™ and 45 µl of lipofectamine3000 were added to another tube. Both tubes were mixed and spun down before combining. The combined mixture was incubated at RT for 15-20 min before dropwise adding into the 10 cm HEK293T plate. Viruses were collected 48 h after transfection, filtered through syringe filters with 0.45 µm pore size (VWR), aliquoted in 1-2 mL and stored at −80 °C. Aliquots were thawed only once to avoid loss of viral viability from repeated freeze–thaw cycles.

### shRNA knockdown in inducible MYC system

To assess MYC-dependent vulnerabilities in ribosome biogenesis, we performed shRNA-mediated knockdown of candidate ribosome biogenesis factors in MCF10A cells expressing inducible MYC through a TetOff system. In this system, MYC overexpression (“MYC-high”) was induced by doxycycline withdrawal, whereas cells maintained in doxycycline were designated “MYC-low.” Unless otherwise indicated, inducible MYC expression was achieved using a TetOff-MYC-HaloTag construct. For experiments in Fig. 1, Fig. 3f, and Extended Data Figs. 1 and 4, a TetOff-MYC-GFP construct was used instead to avoid spectral overlap with endogenously tagged HaloTag-RPL10A and HaloTag-RPS14.

shRNAs targeting ribosome biogenesis factors across multiple processing steps were obtained from Sigma-Aldrich as bacterial glycerol stocks (**Supplementary Table 1**), followed by plasmid preparation and lentiviral packaging. A non-targeting shRNA^31^ (GCTCTTAACTAACGTCACCTA) was used as a negative control. Cells were maintained in MYC-low conditions (200 ng/mL doxycycline) throughout lentiviral transduction in 6-well plates. One day after transduction, shRNA-expressing cells were selected using 2 µg/mL puromycin (Sigma, P4512-1MLX10) under MYC-low conditions. Two independent shRNAs were tested per target when available, to ensure reproducibility and minimize off-target effects; shRNAs that did not achieve efficient knockdown were excluded from further analysis. After puromycin selection, cells were washed extensively (one wash with PBS followed by three 5 min washes with complete media) to remove doxycycline and then split into two conditions: continued culture in doxycycline-containing media (“MYC-low”) or culture in doxycycline-free media (“MYC-high”). shRNA transduction preceded MYC induction to ensure equivalent shRNA exposure and knockdown efficiency between conditions, enabling direct comparison of MYC-dependent cellular responses. Cell proliferation measurements were initiated immediately upon MYC induction and monitored for four days using an Incucyte live-cell imaging system, as described below.

### Live-Cell imaging (Incucyte) to track cell proliferation

Cell proliferation was monitored using the IncuCyte SX5 live-cell imaging system (Sartorius). After shRNA transduction and separation into “MYC-low” and “MYC-high” conditions, cells were seeded into 96-well plates (Fisher Scientific, 167008) at 6,000 cells per well. Plates were imaged every 6 h for four days and cell confluency was quantified to generate growth curves across all knockdown and control groups. Three biological replicates were included per condition.

### Ribosome biogenesis efficiency measurements

Ribosome biogenesis (RiBi) efficiency was defined as the ratio of ribosome production (output) to rRNA transcription (input). Ribosome assembly output was quantified using HaloTag block-labeling to measure the accumulation of newly assembled ribosomes during a defined labeling window, whereas nascent rRNA synthesis input was quantified by short-pulse 5eU incorporation. MCF10A Tetoff-MYC-GFP cells were cultured with or without 200 ng/mL dox for 72 h prior to efficiency measurements.

### Labeling of newly assembled ribosomes through HaloTag Block-Label experiments

To measure the rate of nascent ribosome production, we performed HaloTag Block-Label experiments in endogenously tagged MCF10A cell lines (HaloTag-RPS14 or HaloTag-RPL10A knock-in), as previously described.^35^ Cells were incubated with 25 μM non-fluorescent HaloTag blocking ligand at 37 °C for 30 min to saturate HaloTag binding sites on pre-existing ribosomal subunits. After removal of the blocker (one wash in DPBS followed by three washes with media), cells were incubated with 250 nM HaloTag-TMR ligand (Promega,G8252) for 10 h at 37 °C to label newly assembled ribosomal subunits. TMR signal therefore reports the accumulation of newly assembled ribosomes during the labeling period rather than total ribosome abundance.

For imaging-based quantification, cells were fixed in 4% paraformaldehyde (PFA, 15 min, room temperature) in 96-well glass-bottom plates. Images were acquired and fluorescence intensities quantified as described in *Image Preprocessing, Segmentation, and Quantitative Fluorescence Analysis*. For Western blot-based quantification, cells were harvested on ice in DPBS, followed by cytoplasmic fractionation following established protocols.^70,71^ Briefly, harvested cells were pelleted by centrifuging at 300g for 5 min at 4 °C, resuspended in 150 μL ice-cold 0.3% NP-40 (Sigma Aldrich, 74385) in DPBS supplemented with 1X cOmplete EDTA-free protease inhibitor cocktail and triturated using a p1000 pipette 5 times. The lysate was centrifuged at 16,000g for 30 seconds at 4 °C and the supernatant was collected as the cytoplasmic fraction for subsequent quantification of TMR abundance via Western blotting.

### 5eU pulse labeling to quantify nascent rRNA synthesis (input)

To measure the rate of rRNA transcription, cells were incubated with 1 mM 5eU (Jena Biosciences; CLK-N002-10) at 37 °C for 30 min.

For microscopy-based quantification, cells were fixed in 4% paraformaldehyde (15 min), permeabilized in PBST (PBS + 0.5% Triton X-100, 15 min), and subjected to click chemistry using the Click-iT Plus Alexa Fluor 647 Picolyl Azide kit (Thermo Fisher, C10643) according to the manufacturer’s protocol prior to imaging. Images were acquired and fluorescence intensities quantified as described in *Image Preprocessing, Segmentation, and Quantitative Fluorescence Analysis*.

For dot blot-based quantification, cells were lysed in buffer RLT (QIAGEN, 79216) and total RNA was extracted. Biotin was conjugated to 5eU-containing RNA by click chemistry as previously described.^31,39^ Biotinylated RNA (400 ng per sample) was spotted onto nitrocellulose membranes (Bio-Rad, 1620215), UV-crosslinked, blocked in 0.5% BSA in PBST, and incubated with IRDye 800CW streptavidin (LI-COR, 926-32230; 1:2000 dilution). Membranes were washed and imaged using a LI-COR Odyssey imaging system. Streptavidin signal intensity, proportional to 5eU incorporation, was quantified in Fiji.

### Calculation of RiBi efficiency

RiBi efficiency was calculated after quantification of both assays. For imaging-based measurements, TMR and 5eU signals were quantified in the same cells and RiBi efficiency was computed on a per-cell basis. For biochemical measurements, TMR and 5eU were measured in independent experiments performed under identical culture conditions; within each assay, background-subtracted signals were normalized to MYC-low controls, and RiBi efficiency was calculated as the ratio of the mean normalized TMR signal to the mean normalized 5eU signal.

### Polysome Fractionation and Analysis

Polysome fractionation was performed as described before^31^ with minor modifications. Briefly, cells were lysed in 400 µL Polysome Lysis Buffer (25 mM HEPES, pH 7.3, 150 mM NaCl, 15 mM MgCl₂, 1% Triton X-100, 8% glycerol, 0.5% sodium deoxycholate, 100 μg/mL CHX, 1 mM DTT, RNase inhibitors (NEB, M0314L, 1:60 dilution) and DNase (Thermofisher, AM2239, 1:400 dilution). Lysates were incubated on ice for 15 min, followed by two consecutive centrifugations at 800 x *g* for 5 min, one centrifugation at 8000 x *g* for 5 min, and one centrifugation at 20000 x *g* for 5 min (all at 4 °C) to remove nuclei and mitochondria. RNA concentrations were measured with Qubit RNA BR (Fisher Scientific, Q10211) and lysates were normalized for equal RNA mass before loading onto a 10–50% (w/v) sucrose gradient prepared in Polysome Gradient Buffer (25 mM HEPES, pH 7.3, 150 mM NaCl, 15 mM MgCl₂, 100 μg/mL CHX, and 1 mM DTT).

Ultracentrifugation was performed at 40,000 rpm for 2.5 h at 4 °C in a SW41 Ti rotor (Beckman Coulter). Following ultracentrifugation, gradients were fractionated using a piston gradient fractionator (Biocomp) with continuous monitoring of absorbance at 254 nm to visualize ribosomal profiles. Fractions corresponding to free ribonucleoprotein complexes, 40S and 60S subunits, 80S monosomes, and polysomes (>2 ribosomes) were collected and pooled separately. Protein precipitation from these fractions was performed using the ProteoExtract Protein Precipitation Kit (Fisher Scientific, 53-918-01KIT). After protein quantification with the Pierce Bradford assay (ThermoFisher, 23246), equal amounts of protein were loaded onto Western blotting for quantification of TMR abundance in each fraction.

### 5eU labeling combined with quantitative spike-ins and sequencing

5eU pulse chase RNA-labeling followed by sequencing (5eU-seq)^31^ were performed as previously described with modifications described below. Briefly, RNA lysate was harvested, isolated, click-reacted with biotin picolyl azide (Click chemistry tools, 1167-25), captured using 3 rounds of streptavidin-bead isolation and washes, and sequenced after RNA-seq library preparation

### 5eU pulse-chase labeling in cells

5eU pulse-chase labeling was performed as described.^31^ Cells were seeded in 6 cm dishes a day before 5eU pulse-chase labeling to reach ∼50-60% confluence at the time of harvesting. To label nascent RNA, cells were incubated with 1 mM 5eU (Jena Biosciences; CLK-N002-10) for 15 min at 37 °C with 5% CO_2_. Labeling was terminated by washing cells with 1x DPBS containing 10mM uridine to remove 5eU followed by addition of media with 10 mM uridine (Sigma, U6381) for chasing over various time points. All solutions were kept at 37 °C with a heat block and solution changes were performed as fast as possible at room temperature before putting dishes back into 37 °C incubators to minimize the impact of temperature on RNA transcription and processing in cells. Chases were terminated by aspiration of media and immediate addition of 1X Buffer RLT (QIAGEN, 79216) to lyse the cells. Cell lysates were saved at −80° C before performing RNA isolation.

### In vitro transcription and preparation of ERCC spike-in RNAs

ERCC spike-in RNAs were generated by *in vitro* transcription (IVT) from ERCC^72^ DNA templates (NIST, 2374) containing a T7 promoter. DNA templates were linearized with restriction enzyme BamHI-HF (NEB, R3136S) and gel purified prior to transcription. IVT was performed using the HiScribe T7 RNA Synthesis Kit (NEB, E2040S) at 37 °C for 8–12 h following the manufacturer’s protocol with minor modifications. To generate 5eU–labeled spike-in RNAs, 2% of total UTP was replaced with 5-ethynyl-UTP (5eUTP; Axxora, JBS-CLK-T08-S). Specifically, in a standard reaction containing 2 µL UTP (100 mM), 1.96 µL UTP (100 mM) and 4 µL 5eUTP (1 mM) were used in place of UTP. TURBO™ DNase (ThermoFisher, AM2239) was used to remove the template DNA after IVT. ERCC RNAs were purified with the Zymo RNA Clean and Concentrator-25 kit (R1017), quantified with Qubit RNA BR (Fisher Scientific, Q10211) and assessed for size and integrity on 2% E-Gel™ EX Agarose Gels (ThermoFisher, G402022). Purified RNAs were pooled at equal-molar ratio and stored at −80 °C. Selected spike-in RNA sequences are listed in Supplementary Table 1.

### ERCC spike-in addition and validation

IVT ERCC RNAs were added to each RNA sample at a fixed amount proportional to the total RNA mass (0.3 ng of spike-in RNAs per 12 μg of total RNA) to enable normalization across samples. ERCC spike-ins were added prior to click chemistry so that 5eUTP-containing spike-ins underwent identical handling, purification, and library preparation steps as cellular 5eU-labeled RNA. To test if the 5eU-seq spike-ins could quantitatively measure the amount of 5eU-labelled RNA in a sample, mock RNA samples with 0X, 1X, 2X, 4X and 6X of 5eU-labelled total RNA were prepared by mixing 5eU-labeled total RNA samples with non-labeled total RNA samples at different ratios (**Extended Data Fig. 3a-b**). Spike-in RNAs were added, followed by click reaction and capture of 5eU-labeled RNA (as described below). RT-qPCR was used to determine the ratios of captured 5eU cellular RNAs to spike-in RNAs. The measured results were plotted against the expected ratios (**Extended Data Fig. 3c-d**).

### Biotinylation, capture, and RNA-seq library preparation of 5eU-labeled RNA

To biotinylate 5eU-labeled RNA, click chemistry was performed on spike-in-added total RNA (10-20 µg) isolated from 5eU-pulse chase following previous protocols.^31,39^ Biotinylated RNA was then captured with Dynabeads MyOne Streptavidin C1 beads (Invitrogen, 65002) with three rounds of captures and washes as previously described to efficiently remove background from mature rRNAs.^31^ 5eU-seq libraries were prepared^73^ and sequenced on a NovaSeq (Illumina) with paired-end reads (150×150).

#### Mammary tumor xenograft model

All animal procedures were performed in accordance with protocols approved by the Institutional Animal Care and Use Committee (IACUC) of Princeton University. Mice were maintained in a controlled environment at 20–22 °C with a 14 h light / 10 h dark cycle and 40–70% relative humidity. MCF10CA or SUM159 cells expressing control shRNA or two independent shRNAs targeting PES1 (TRCN0000117722 and TRCN0000300648) were injected into the left fourth mammary fat pad of female NOD/SCID Gamma (NSG) mice (8-12 weeks old) at a density of 1×10⁵ cells in 10 µl. Tumor growth was monitored weekly. Tumor length and width were measured with calipers, and tumor volume was calculated as (length x width^2)/2. Mice were sacrificed 7 weeks after injection, and tumors were excised and weighed.

#### TCGA survival analysis (Kaplan-Meier curves)

Gene expression and clinical data for TCGA primary tumors were obtained from the UCSC Xena browser (TCGA Pan-Cancer). Only samples annotated as primary tumors were included. MYC expression and ribosome biogenesis (RiBi) factor expression scores were calculated from normalized RNA-seq expression values. Guided by cryo-EM structural characterization of ribosome assembly intermediates,^11,50^ we curated gene sets representing early pre-rRNA processing (co-transcriptional cleavage at sites 01/02) and late processing (ITS2 maturation at sites 3′/4′ and cytoplasmic maturation at site 3) (**Supplementary Table 1**). Patients were stratified into MYC-high and MYC-low groups using the median MYC expression across the cohort to define an unbiased, outcome-independent threshold. Within each MYC stratum, tumors were further classified as RiBi-high or RiBi-low based on the median expression of the corresponding early or late ribosome biogenesis (RiBi) gene set. Overall survival time and event status were obtained from UCSC Xena clinical annotations. Follow-up times were truncated at 3000 days to reduce the influence of sparse long-term observations and stabilize hazard estimation. Survival curves were estimated using the Kaplan–Meier method and compared using two-sided log-rank tests. To determine whether the prognostic effect of RiBi expression depended on MYC status, multivariable Cox proportional hazards models were fit including MYC group, RiBi group, and a MYC×RiBi interaction term. Hazard ratios (HRs) were calculated from regression coefficients (HR = e^β), and statistical significance was assessed using two-sided Wald tests. All analyses were performed in R (survival and survminer packages).

#### Image Preprocessing, Segmentation, and Quantitative Fluorescence Analysis

Raw microscopy image files (.nd) were acquired as multi-channel three-dimensional z-stacks and exported as TIFF stacks using Nikon Elements (Nikon Instruments Inc., Tokyo, Japan). To reduce computational burden while preserving cellular morphology and intensity distributions, all z-stack images were downsampled by a factor of four in the x–y dimensions using ImageJ/Fiji^74^ (National Institutes of Health, Bethesda, MD, USA). Image stacks were visually inspected to ensure that downsampling did not compromise segmentation fidelity. A custom ImageJ macro was developed to automate channel separation, z-stack organization, and metadata preservation across all datasets. Following channel splitting, image stacks were assigned as Channel 1 (C1): 647, nascent transcripts (5eU) labeling; Channel 2 (C2): 568, nascent ribosome (TMR) labeling, Channel 3 (C3, not used in analyses): 488, steady-state ribosome (Oregon Green) labeling; and Channel 4 (C4): 405, DAPI. Channel alignment and consistency across z-slices were verified prior to downstream analysis.

Three-dimensional whole-cell segmentation was performed using the C2 channel, which provided robust contrast for delineating cell boundaries. Segmentation was carried out using Cellpose^75^ (Howard Hughes Medical Institute, Ashburn, VA, USA) with the pretrained *cyto2_cp3* model. Model performance was evaluated by visual inspection of segmentation masks across multiple fields of view and z-slices, and segmentation outputs were curated to exclude partial cells at image boundaries. Nuclear segmentation was performed independently in three dimensions using the C4 channel, which selectively labeled nuclei. The pretrained *nuclei* model in Cellpose was applied to generate 3D nuclear masks. Nuclear segmentation accuracy was validated by overlaying masks on raw fluorescence images to ensure correct identification of nuclear volumes and to avoid over-segmentation or merging artifacts.

Following segmentation, z-stack images from channels C1, C2, and C4, together with the corresponding 3D whole-cell and nuclear masks, were imported into CellProfiler^76^ (Broad Institute, Cambridge, MA, USA) for quantitative analysis. A custom CellProfiler pipeline was constructed to associate each nucleus with its corresponding whole cell, propagate nuclear and cellular labels through the full z-stack, segment nucleoli from channel C1, and extract fluorescence intensity measurements on a per-cell basis. Background intensity was measured from cell-free regions of images and subtracted from all intensity measurements. Fluorescence intensity of C1 and C2 was quantified within the three-dimensional whole-cell, nuclear, and nucleolar regions. Integrated fluorescence intensity was computed by summing voxel intensities across all z-slices for each compartment. Cytoplasmic ribosome fluorescence intensity was calculated by subtracting nuclear intensity from the corresponding whole-cell intensity for each individual cell, thereby isolating the cytoplasmic signal while accounting for cell-to-cell variability in expression and size.

Quantitative outputs from CellProfiler were exported and further processed using custom scripts written in MATLAB (MathWorks, Natick, MA, USA). These scripts were used to calculate integrated and compartment-specific fluorescence metrics and aggregate single-cell measurements across experimental conditions. Final datasets were compiled for downstream statistical analysis and visualization in GraphPad Prism 10.

### Computational analysis of 5eU-seq, spike-in normalization and 2’-*O*-Methylation

#### 5eU-seq with internal spike-in control to quantify cleavage rates and abundance

5eU sequencing reads were aligned and quantified using the previously reported workflow^31^ with modifications to support alignment to multiple genomes with STAR (v2.7.11): https://github.com/SoftLivingMatter/5eu-seq-pipelines. After adaptor and quality trimming, reads were aligned to multiple reference genomes independently and in parallel by default. To quantify the number of reads aligning to different regions in pre-rRNA, mRNAs, or spike-in RNAs, a new ‘region count’ analysis was introduced, which counts the total number of reads aligning to different genomic positions (regions of interest) from a bed file using bedtools (v2.24) intersect. To normalize read counts and estimate relative RNA abundance, orthogonal spike-in sequences were added to samples in known quantities. In the workflow, a genome for the spike-in sequences was provided along with regions encompassing the entire spike-in sequence. For normalizing read counts, outputs of counts for each region of interest (e.g., mRNAs, pre-rRNAs) were then normalized by the reads present in the spike-in region.

Cleavage rates at all sites were measured by quantifying the fraction of non-spanning reads (“cleaved”) relative to total reads (“cleaved” + “uncleaved”). The positions of these sites on human rDNA, along with the quantification windows, were previously described.^31^

Note that a correction was applied when quantifying spike-in–normalized counts for pre–site 3 (uncleaved) and post–site 3 (cleaved) rRNA species. The site 3 cleavage site lies in close proximity to the conserved dimethylation site at the 3′ terminal loop of 18S rRNA,^77–79^ which induces reverse transcription (RT) stops and results in the loss of short 18S 3′-terminal fragments. Consequently, the absolute read counts for both cleaved and uncleaved species at site 3 are substantially underestimated due to interference from this modification. To correct for this technical loss, we quantified total read coverage across a 100-nt internal region of the 18S rRNA (∼500 nt upstream of the site 3 cleavage site) that is not affected by RT termination. Read counts at site 3 were then scaled to match the coverage of this internal reference region, assuming that in the absence of RT termination artifacts, the 18S 3′ end would exhibit the same decay kinetics as the unaffected internal region. Cleaved and uncleaved species were scaled proportionally.

### 2’-O-methylation analyses

2’-*O*-methylation levels on 18S and 28S rRNA were quantified over time from 5eU-seq data using the previously reported workflow^31^ adopted from RiboMethSeq.^80,81^ “Early” and “Late” 2’-*O*-methylation sites were defined based on the time it takes to reach 90% of the maximal modification level (t_90_) in MCF10A cells. Modification sites with average t_90_ (n = 2 replicates) value equal to or less than 60 min are classified as “early” sites while sites with t_90_ more than 60 min are classified as “late” sites. Modification sites with high variations between replicates (s.e.m > 15 min) were challenging to define modification rates for and thus excluded from the analysis. A table of high-confidence early and late 18S and 28S rRNA 2’-O-methylation sites can be found in Supplementary Table 1. The average of all high-confidence early sites or late sites were shown per condition.

#### Estimation of the delay time from fraction cleaved measurements

In Fig. 2 of the main text, we defined the delay time Δτ ≡ τ*_malignant_* – τ*_normal_* as the difference between the average time to cleave a site in malignant versus normal cells. For a given site, let *f*(*t*) be the fraction of pre-rRNA with that site cleaved at time *t*, with 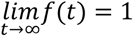. The average cleavage time is given by 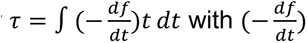 representing the fraction of cleavage that takes place at time *t*. The delay time is thus given by:

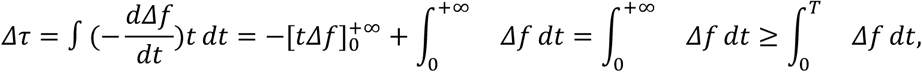

where *T* is the duration of the experiment. Thus, the delay time can be lower bounded by the area between the fraction-cleaved curves (Fig. 2c).

To estimate the delay time Δτ from 5eU-seq measured cleavage fractions, we use a trapezoidal scheme:

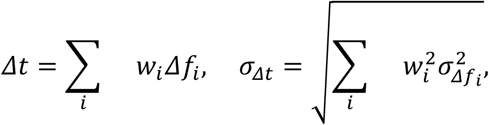

where Δ*f_i_* = *f_i,normal_* – *f_i,malignant_* is the difference in the fraction cleaved at time point *i*, with σ_Δ*fi*_ being the corresponding experimental uncertainty. The trapezoidal weights are given by: 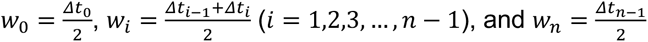, with Δ*t_i_* = *t_i_*_+1_ – *t_i_*.

## Data availability

Data supporting these findings will be made available on public databases (GEO) upon publication or upon request.

## Code availability

All code and pipelines used for this study are provided in the following repositories: https://github.com/SoftLivingMatter/5eu-seq-pipelines.

## Materials and correspondence

Correspondence and requests for materials should be addressed to Clifford P. Brangwynne.

## Acknowledgements

We thank all Brangwynne lab and Kang lab members and E. Soehalim for experimental help and helpful discussions; A. Zhu and M. Iglesias of the Wuhr laboratory for assistance with sucrose gradient fractionation; D. Sanders and A. Lin for gifted plasmids; P. Bhat and M. Guttman for 5eU-seq assistance; N. Jaberi-Lashkari, N. Patel, M. Ebert, M. Lee, and A. Donic for manuscript feedback; E. Gatzogiannis and the Molecular Biology Confocal Imaging Facility for microscopy assistance; C. DeCoste, K. Rittenbach, G. Palmieri and J. Garcia and the Molecular Biology Flow Cytometry Resource Facility, which is partially supported by the Rutgers Cancer Institute of New Jersey NCI-CCSG P30CA072720-5921, for FACS cell sorting support; W. Wang and the Genomics Core Facility for library preparation and sequencing.

This work was supported by the Howard Hughes Medical Institute, the Princeton Biomolecular Condensate Program, the Princeton Center for Complex Materials, a MRSEC (NSF DMR-2011750), the St. Jude Collaborative on Membraneless Organelles, the AFOSR MURI (FA9550-20-1-0241), the Chan Zuckerberg Initiative Exploratory Cell Network, Breast Cancer Research Foundation, Ludwig Cancer Research, Brewster Foundation, American Cancer Society and Susan G. Komen Foundation. Q.Y. is supported by the Harold W. Dodds Fellowship from Princeton University. S.A.Q. is supported by an HHMI Hanna H. Gray Fellowship. J.F.B. is supported by the NSF GRFP Fellowship.

## Author Contributions

L.J., Y.K., and C.P.B. designed the study. Q.Y., and A.K. performed mathematical modeling. L.J., S.A.Q., J.F.B. S.A., A.A.A. and J.T. performed experiments. L.J., S.A.Q., A.A.A. and T.C. performed genomic data analyses. J.X. performed image analyses. L.J. and Y.W. performed clinical patient data analyses. L.J., Q.Y., and C.P.B. wrote the manuscript with input from all authors. L.J. and Q.Y. made the figures with contributions from all authors. C.P.B. and Y.K. supervised the project.

## Supplementary Information

**Supplementary Note 1**

Pulse-chase labeling with 5-ethynyl-uridine (5eU) was used to measure pre-rRNA dynamics over time. In an ideal pulse-chase experiment, incorporation of the labeled nucleoside would stop immediately upon addition of excess unlabeled uridine during chase. In practice, intracellular nucleotide pools do not equilibrate instantaneously. As a result, a small fraction of newly synthesized RNA can continue to incorporate 5eU for a short period after the start of the chase. To account for this, we incorporated labeling kinetics into the mathematical model (See Supplementary Note 2).

**Supplementary Note 2 (Attached)**

Mathematical modeling of pre-rRNA synthesis, cleavage, and degradation.

**Supplementary Note 3**

To estimate ribosome biogenesis efficiency from the model, we used the fraction of nascent transcripts remaining at the final cytoplasmic cleavage step in the SSU pathway (site 3) as a proxy for ribosome yield. This step occurs late during maturation of pre-40S particles and marks the final rRNA processing event before formation of mature 18S rRNA and entry into the functional ribosome pool. By this stage, upstream assembly events, including rRNA folding and incorporation of ribosomal proteins, have largely occurred, so the fraction of transcripts reaching site 3 reflects the proportion of nascent pre-rRNAs that successfully complete ribosome assembly.^82–84^ In contrast, the final LSU cleavage events monitored in our 5eU-seq analysis (sites 3′ and 4′) occur on nuclear pre-60S particles before completion of late maturation events, including structural remodeling, maturation of the 5S RNP-containing central protuberance, and subsequent nuclear export and cytoplasmic quality-control steps.^11,83,85^ Efficiencies derived from these LSU cleavages therefore do not capture losses that occur during these later stages of ribosome assembly. For this reason, we used site 3 as a more comprehensive estimator of ribosome biogenesis efficiency in our model.

**Supplementary Note 4**

Direct experimental measurement of ribosome biogenesis efficiency using endogenous ribosomal protein tagging was not feasible in MCF10CA (malignant) cells. This assay requires generation of clonal cell lines carrying endogenously tagged ribosomal proteins (with HaloTag) to enable pulse-labeling of newly assembled ribosomes. However, repeated attempts to generate such tagged clones in MCF10CA cells resulted in substantial alterations in cellular fitness and deviations from the parental malignant phenotype. Establishing these clones requires prolonged single-cell clonal expansion from an individual edited cell, a process that can take weeks to months and introduces substantial phenotypic drift in MCF10CA cells. Because these clonal effects confounded interpretation of ribosome production measurements, we did not perform TMR-based efficiency measurements in this system and instead relied on model-derived estimates of ribosome biogenesis efficiency.

**Supplementary Note 5**

Choice of system for perturbation experiments: shRNA knockdown experiments were performed in the inducible MYC system for both ribosome biogenesis measurements and proliferation assays. In this system, cells were transduced with shRNA and selected with puromycin prior to MYC induction, so MYC-low and MYC-high conditions experienced identical viral exposure, selection history, and intracellular shRNA levels. Upon induction, the only experimental variable is MYC activation, allowing perturbation strength to be matched across conditions. In the MCF10A/MCF10CA pair, comparable control is not feasible. The two lines differ in proliferation rate, transduction efficiency, and selection tolerance, leading to unequal intracellular shRNA dosage even under identical infection conditions. Because ribosome biogenesis output and cell growth are highly sensitive to gene dosage, differences in knockdown efficiency would be confounded with differences in oncogenic state. We therefore used the inducible MYC system to ensure that functional effects on ribosome biogenesis and growth could be interpreted without variability in perturbation magnitude.

**Supplementary Table 1 (Attached)**

Primers, antibodies, plasmids, spike-in sequences, and classification of rRNA modifications and ribosome biogenesis factors by processing stage used in this study.

---

**Extended Data Fig. 1:**
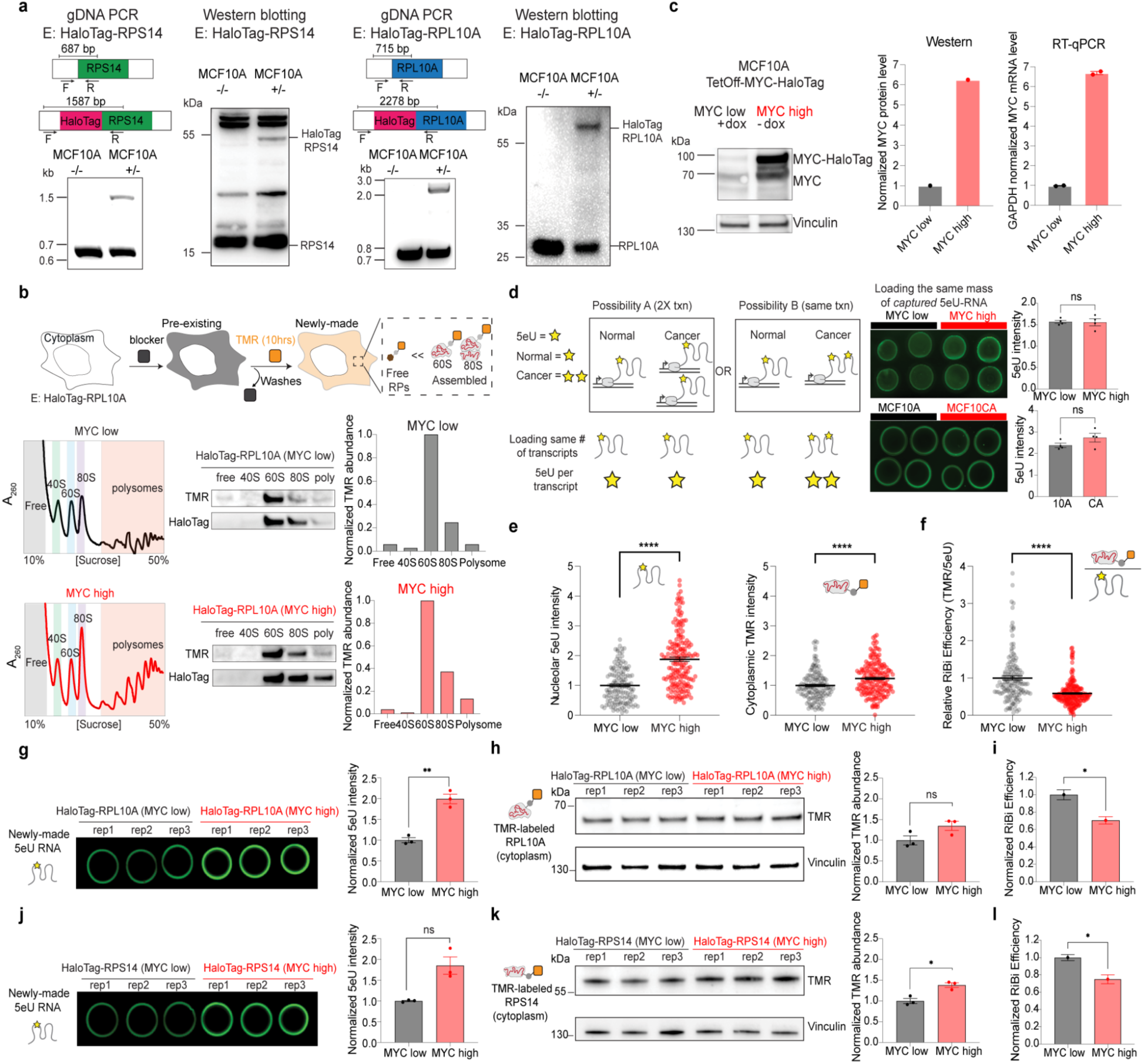
Validation of cell lines and approaches used for ribosome biogenesis efficiency measurements. **a**, Validation of endogenous HaloTag knock-in at the RPS14 or RPL10A loci in MCF10A cells (labeled as E: HaloTag-RPS14; E: HaloTag-RPL10A). Genomic DNA PCR with primers flanking the insertion site confirms integration of the HaloTag sequence upstream of the RPS14 or RPL10A coding regions (MCF10A +/−). Immunoblotting verifies expression of HaloTag-fused RPS14 and RPL10A proteins. **b**, HaloTag-RPL10A cells were blocked and pulse-labeled with TMR ligand for 10 hours to mark newly synthesized RPL10A. Sucrose-gradient fractionation separates free, 40S, 60S, 80S, and polysome fractions, followed by western blotting for TMR and HaloTag. Quantification shows that most TMR-labeled RPL10A localizes to 60S and 80S fractions rather than the free fraction, indicating TMR signal primarily reflects nascent ribosome production. **c**, MCF10A cells expressing the TetOff-MYC-Halotag system. Western blot confirms doxycycline-regulated MYC-HaloTag expression: MYC-low (+ Dox) and MYC-high (- Dox) after 72 h of MYC induction. Quantification of MYC protein (normalized to vinculin) and MYC mRNA levels (RT-qPCR normalized to GAPDH) verifies robust MYC upregulation upon doxycycline withdrawal. **d**, Increased total 5eU signal in oncogenic conditions could reflect either increased transcription (trxn, Possibility A) or increased labeling per transcript (Possibility B). When equal numbers of captured 5eU-labeled transcripts are loaded, 5eU intensity is comparable between MYC states and between MCF10A and MCF10CA cells, indicating that oncogene activation primarily increases transcript abundance rather than incorporation efficiency (n = 4 per condition); two-tailed Welch’s t-test. **e**, MCF10A cells (E: HaloTag-RPS14) are labeled with both 5eU and TMR in MYC-low and MYC-high conditions. Left, quantified 5eU intensity in the nucleolus for rRNA transcription rate. Right, quantified TMR intensity in the cytoplasm for nascent ribosome production rate. **** p value <0.0001 (two-tailed Mann-Whitney test); n = 179 (MYC-high), 140 (MYC-low) cells. **f**, Distribution of relative RiBi efficiency (TMR/5eU) in MYC-low and MYC-high cells, quantified from e, in HaloTag-RPS14 cells. ****p < 0.0001 (two-tailed Mann-Whitney test); n = 179 (MYC-high), 140 (MYC-low) cells. Black lines denote mean values; error bars are s.e.m, in both e and f. **g-l**, Bulk quantification of ribosome biogenesis efficiency in HaloTag-RPL10A cells (g-i) and HaloTag-RPS14 cells (j-l). g, j, Quantification of 5eU incorporation via dot blot after a 30-min 5eU pulse; equal amounts of total RNA were loaded across conditions. h, k, Quantification of cytoplasmic TMR signal by western blot following cytoplasmic fractionation (30-min block and 10-hour TMR labeling). i, l, Ribosome biogenesis (RiBi) efficiency, calculated as TMR/5eU and normalized to MYC-low conditions. Statistical significance was assessed using Welch’s t test: g, ** p value = 0.0045; i, * p value = 0.017; k, * p value = 0.0104; l, * p value = 0.0210. n=3 per condition. Error bars are s.e.m.

**Extended Data Fig. 2:**
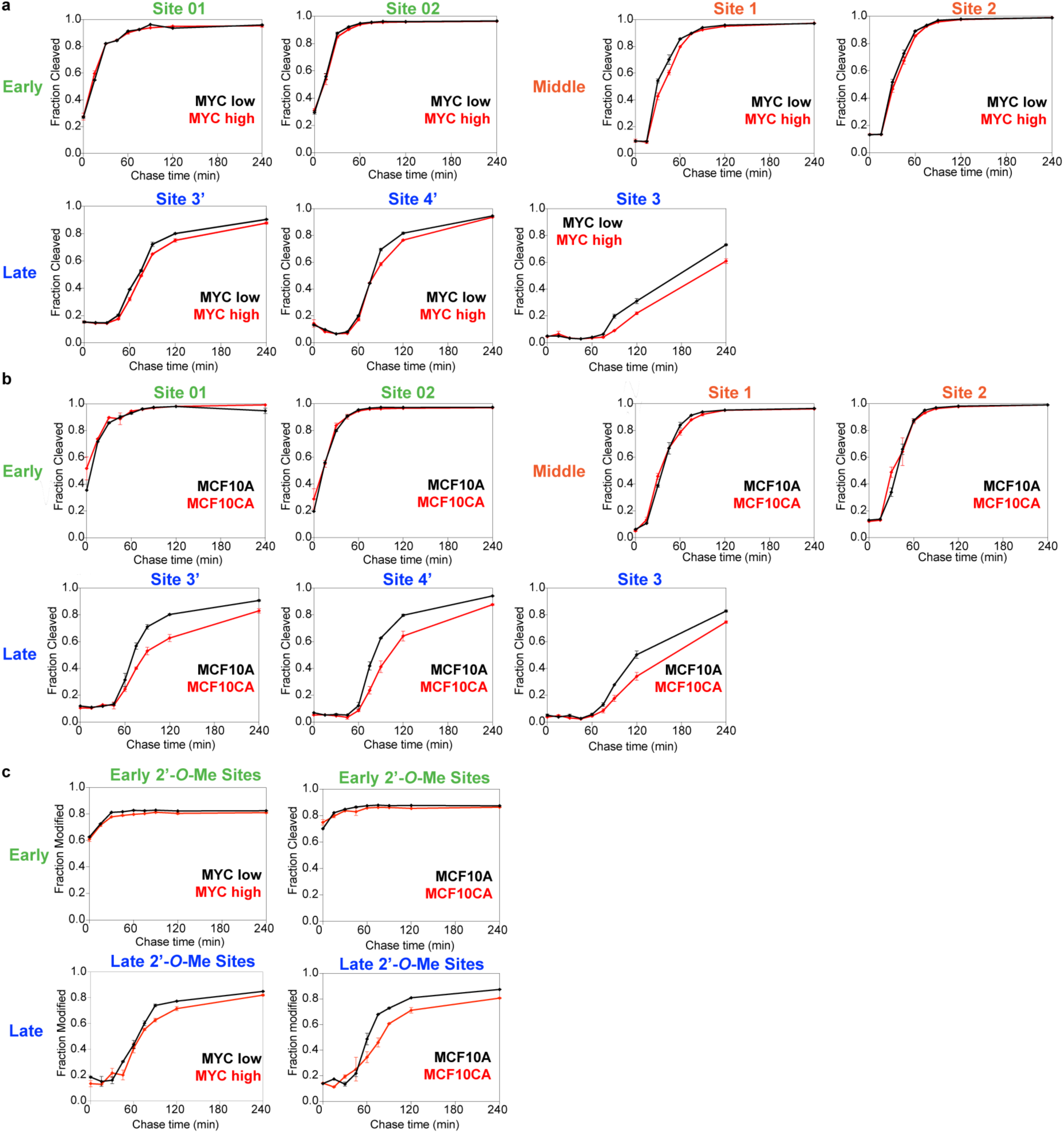
**Pre-rRNA cleavage and modification kinetics a-b**, Fraction of pre-rRNAs cleaved over time at early (green), middle (orange), and late (blue) cleavage sites comparing MYC-low and MYC-high states (a) and MCF10A and MCF10CA cells (b). n = 2 per time point. Black lines denote MYC-low or MCF10A conditions and red lines denote MYC-high or MCF10CA conditions. **c**, Fraction of pre-rRNAs modified at early and late 2′-*O*-methylation sites over chase time in MYC-low versus MYC-high states (left) and MCF10A versus MCF10CA cells (right). n = 2 per time point. See Supplementary Table 1 for classification of early and late modification sites. Error bars are s.e.m in a-c.

**Extended Data Fig. 3:**
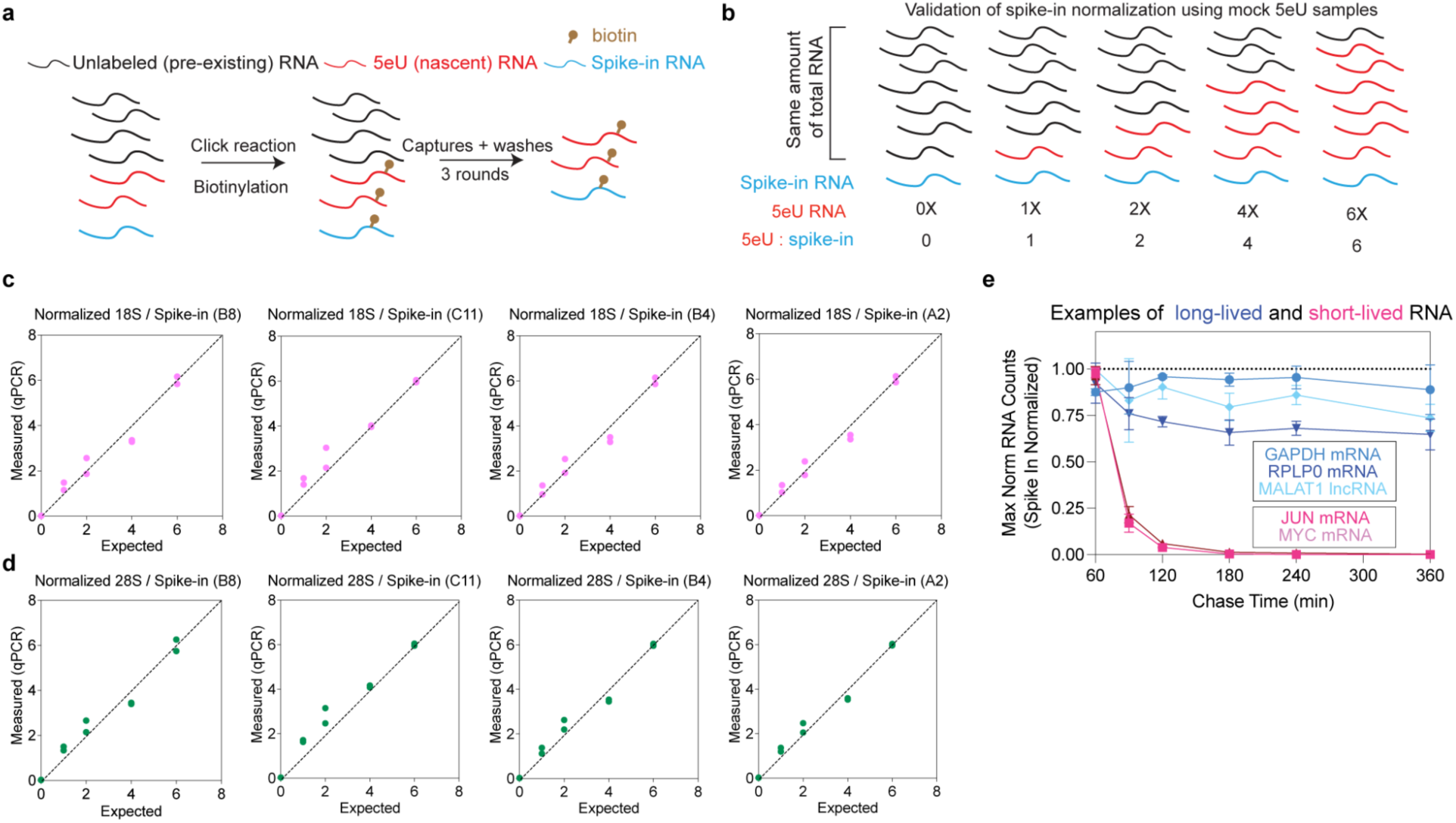
Validation of spike-in-normalized 5eU-seq quantification. **a**, Pre-existing (unlabeled) RNA, nascent (5eU-labeled) RNA, and exogenous spike-in RNA are subjected to click-chemistry biotinylation followed by streptavidin capture and washes prior to downstream quantification. **b**, Design of mock 5eU samples for validation. Schematic illustrating RNA mixtures containing increasing amounts of 5eU-labeled RNA relative to total RNA (0X-6X) with a constant amount of spike-in RNA. **c-d**, Expected (x-axis) versus measured (y-axis) abundances of spike-in-normalized 18S- (c) or 28S- (d) containing pre-rRNAs determined by qPCR. Multiple spike-in sequences (B8, C11, B4, A2) were used. The measured rRNA:spike-in ratios were linearly scaled so that the mean value in the 6X condition was set to 6. *n* = 2 replicates per condition. **e**, Spike-in-normalized abundance of representative cellular long-lived (GAPDH, RPLP0, MALAT1) and short-lived (JUN, MYC) RNAs measured over chase time following 5eU labeling.

**Extended Data Fig. 4:**
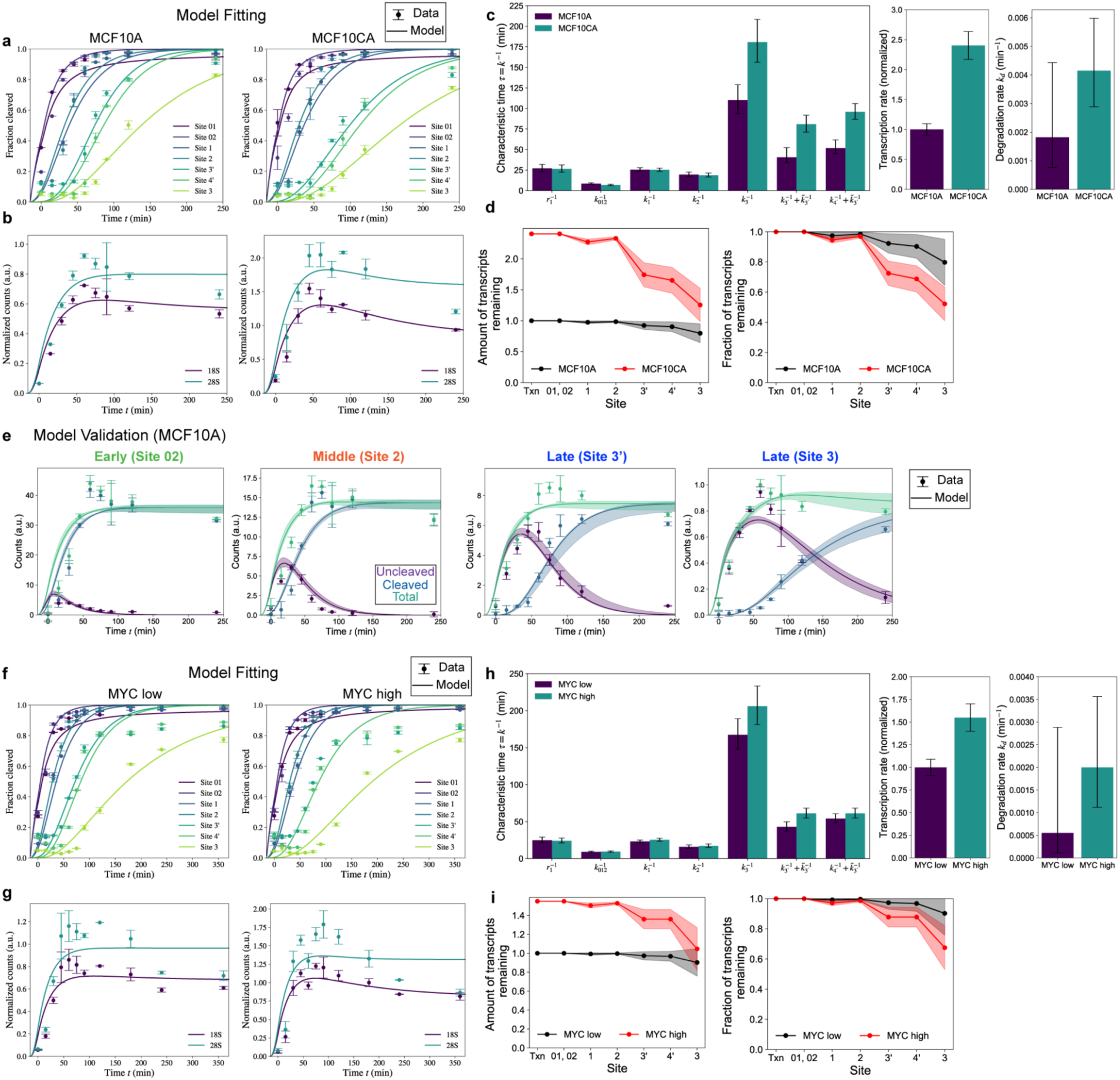
**Validation of the mathematical model and inferred rate parameters a-b**, Model fitting results for MCF10A and MCF10CA. The model was fitted to the fraction cleaved for each cleavage site (a), and the total counts of 18S- or 28S-containing pre-rRNAs (b). **c**, Best-fit cleavage, transcription and degradation parameters from the mathematical model in MCF10A/MCF10CA cells, see Supplementary Note 2 for details. **d**, Model-derived total amount of nascent transcripts remaining (left) and fraction of transcripts remaining (right, normalized to transcription) along the rRNA processing pathway in MCF10A and MCF10CA cells. **e**, Model validation (with data not used for fitting) in MCF10A cells: Model-predicted dynamics (lines) of cleaved, uncleaved and total (cleaved + uncleaved) species at each cleavage step compared with independent experimental spike-in-normalized measurements (points, mean ± s.d., n = 2 per time point); see Supplementary Note 2 for validation in all cleavage sites. **f-g**, Model fitting results for MYC-low and MYC--high cells. The model was fitted to the fraction cleaved (f) and the total counts of 18S- or 28S-containing pre-rRNAs (g). **h**, Best-fit cleavage, transcription and degradation parameters from the mathematical model in MYC-high and MYC-low cells; see Supplementary Note 2 for details. **i**, Model-derived total amount of transcripts remaining (left) and fraction of transcripts remaining (right, normalized to transcription) along the rRNA processing pathway in MYC-low and MYC-high cells. In d, e and i, shaded regions indicate uncertainty from model fitting (see Supplementary Note 2).

**Extended Data Fig. 5:**
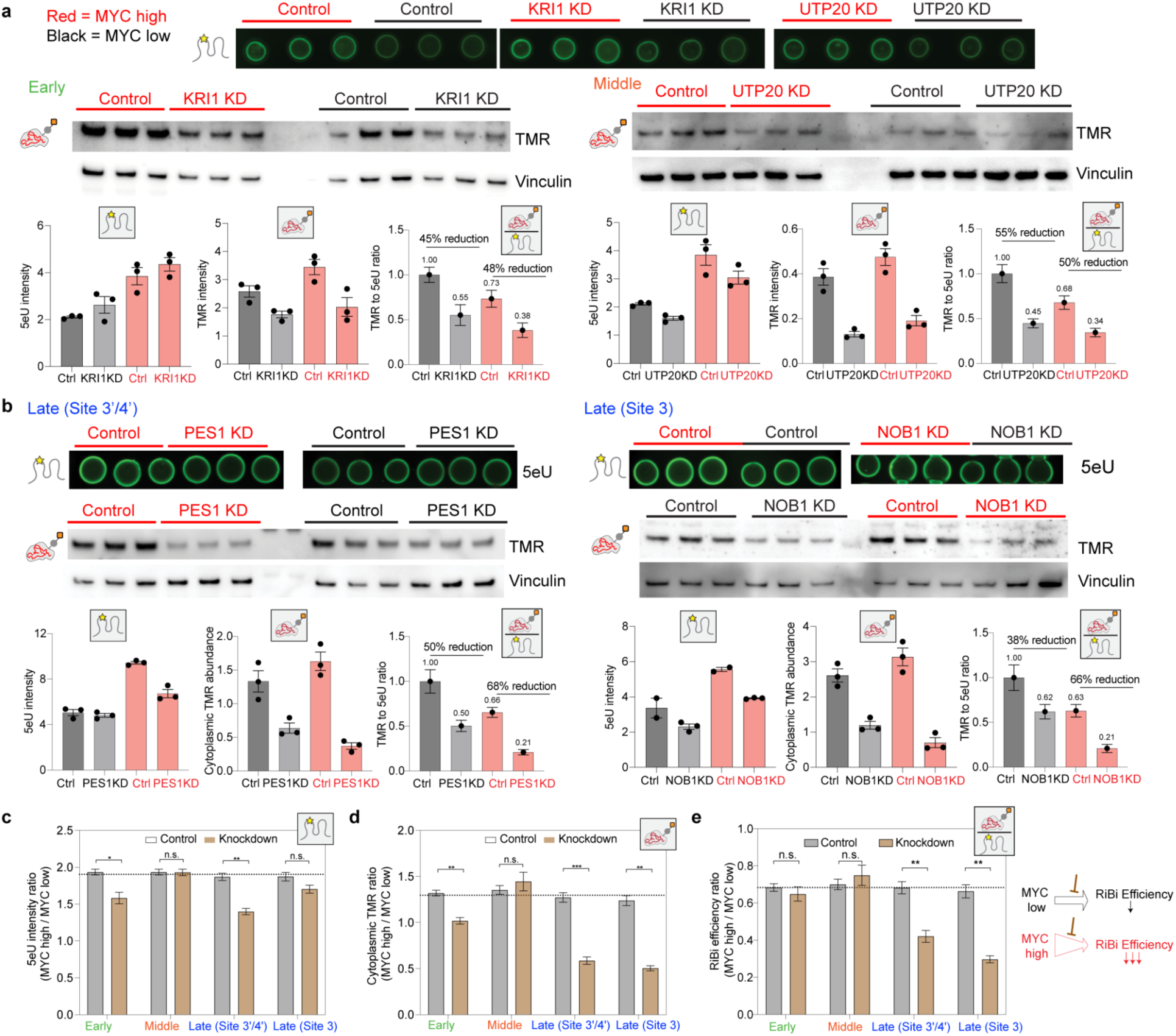
**Late-stage perturbations selectively reduce ribosome biogenesis efficiency in MYC-high cells a-b**, Representative 5eU dot blots (top) and cytoplasmic TMR-Halo ribosome western blots (middle) following knockdown (KD) of early (KRI1, a), middle (UTP20, a) rRNA, or late maturation factors (PES1 and NOB1, b) in MYC-high and MYC-low cells. Red labels denote MYC-high cells and black labels denote MYC-low cells. Vinculin serves as a loading control for TMR blots. Halotag-RPL10A cells were used for PES1 (LSU) KD while Halotag-RPS14 cells were used for KRI1, UTP20 and NOB1 (SSU) KDs. Quantification of 5eU signal intensity, cytoplasmic TMR signal, and TMR-to-5eU ratios are shown below each panel. Bar plots represent mean ± s.e.m. (n = 3 biological replicates). **c**, Quantification of nascent rRNA synthesis from 5eU dot blots in a-b. Y-axis, 5eU signal intensity in MYC-high relative to MYC-low cells. The dotted line indicates the mean MYC-high to MYC-low 5eU ratio across controls; *p = 0.0292 (Early); **p = 0.0022 (Late, Site 3’/4’); two-tailed Welch’s t-test. **d**, Quantification of cytoplasmic ribosome production. Processing factors acting at early (KRI1), middle (UTP20), or late (PES1 and NOB1) steps were knocked down in MYC-low and MYC-high cells. Y-axis, cytoplasmic TMR-Halo signal in MYC-high relative to MYC-low cells. The dotted line indicates the mean MYC-high to MYC-low TMR ratio across controls; **p = 0.0034 (Early); ***p = 0.0006 (Late, Site 3’/4’); **p = 0.0013 (Late, Site 3); two-tailed Welch’s t-test. Error bars denote the propagated s.e.m. (n = 3 per condition) in c-d. **e**, Quantification of ribosome biogenesis (RiBi) efficiency (TMR/5eU) following knocking down processing factors acting at early, middle or late steps in MYC-low and MYC-high cells. Y-axis, ratio of RiBi efficiency in MYC-high relative to MYC-low cells. The dotted line indicates the mean MYC-high to MYC-low efficiency ratio in controls; values below this line indicate a stronger reduction of RiBi efficiency in MYC-high cells. Error bars denote s.e.m., calculated from the bootstrap distribution obtained by resampling biological replicate (n = 3 per condition). **p = 0.0047 (Site 3’/4’); **p = 0.0027 (Site 3); two-tailed Welch’s t-test.

**Extended Data Fig. 6:**
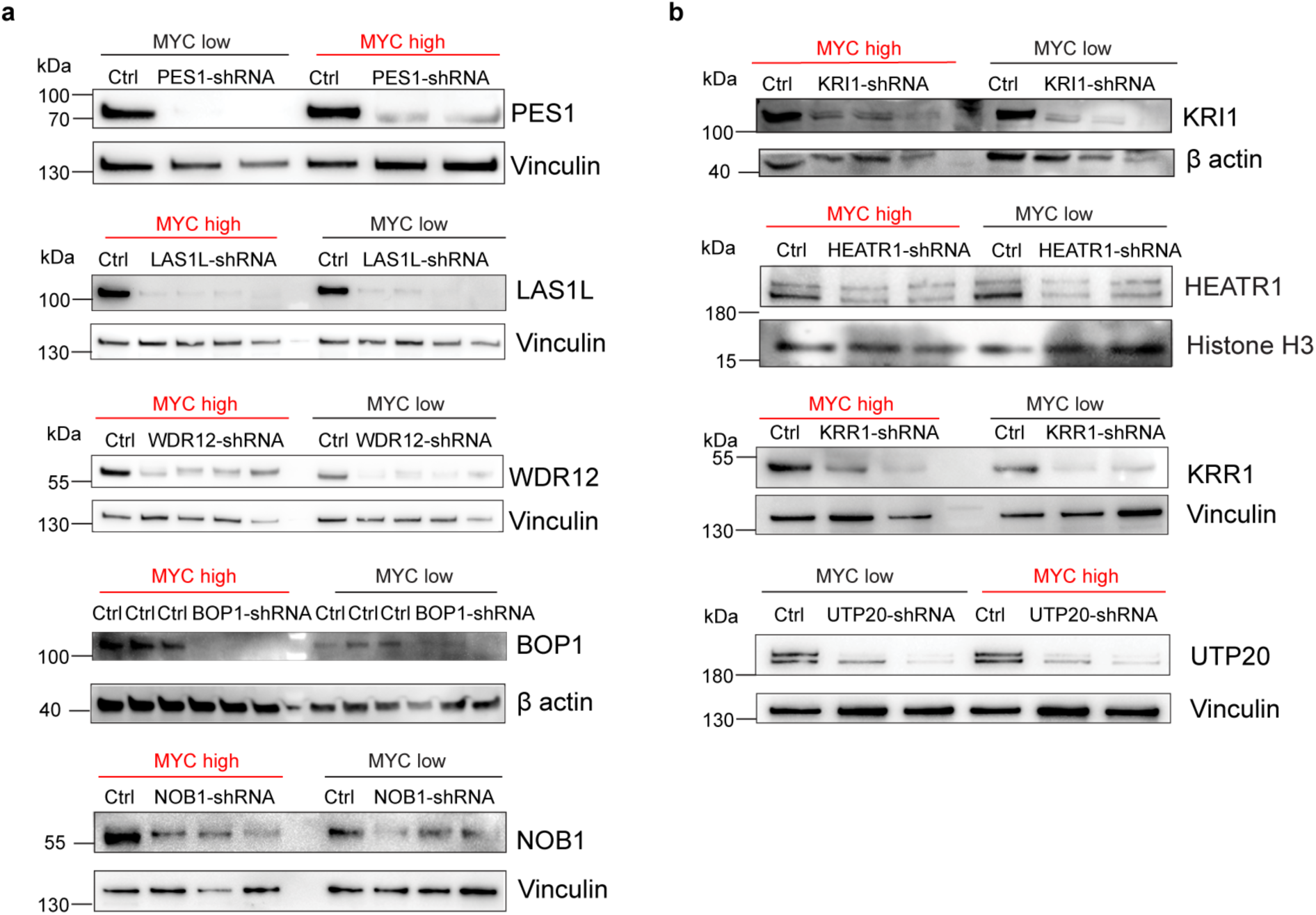
**Western blot validation of ribosome biogenesis factor knockdowns a-b**, Western blotting confirming knockdown of late (a) and early/middle (b) ribosome biogenesis factors in MYC-low and MYC-high cells, for perturbations analyzed in Fig. 4 and Extended Data Fig. 5. UTP15 is not shown since available antibodies did not yield a specific immunoblot signal.

**Extended Data Fig. 7:**
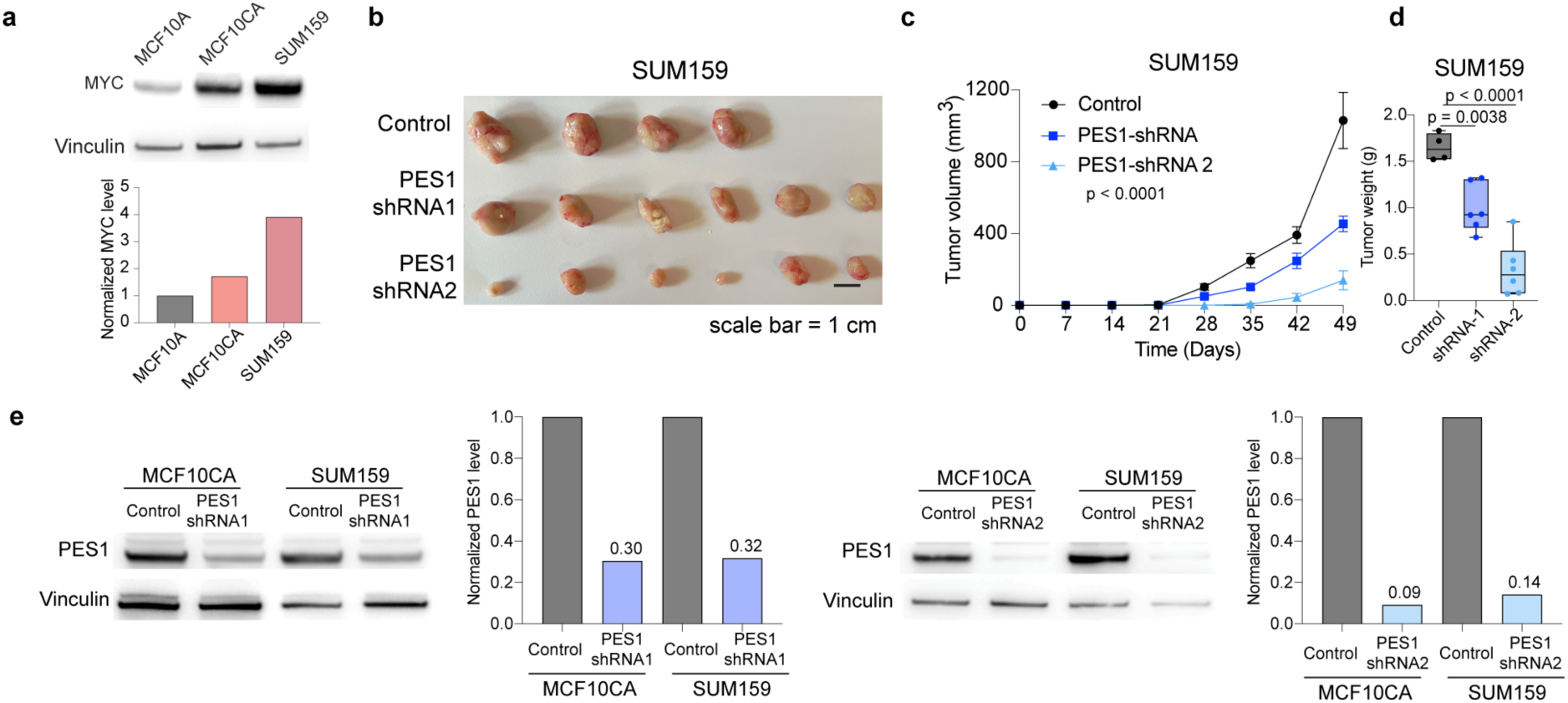
MYC expression and PES1 knockdown in cell lines used for tumor xenografts. **a**, Left, Western blotting of MYC in MCF10A, MCF10CA, and SUM159 cells with vinculin as a loading control. Right, quantification of MYC protein intensity normalized to vinculin. **b-d**, Growth of SUM159 xenograft tumors following PES1 knockdown. b, Representative images of endpoint tumors excised from female NSG mice injected with control cells or cells expressing PES1 shRNAs (shRNA1 or shRNA2). Scale bar = 1 cm. c, Tumor volume over time. Error bars represent mean ± s.e.m. Tumor growth curves were compared using two-way repeated-measures (RM) ANOVA with Geisser-Greenhouse correction; P values indicate the time x treatment interaction. d, Tumor weight at endpoint. Box plots show medians (lines), boxes (25th-75th percentiles), and whiskers (min-max). Statistical significance was assessed using one-way ANOVA with Tukey’s test for multiple comparisons. n = 5 (SUM159 control); 6 (SUM159 PES1-shRNA1); 6 (SUM159 PES1-shRNA2) mice for tumor growth curves. One mouse from the control group died before endpoint tumor collection and was therefore excluded from tumor weight measurements. Final tumor weight analyses were performed with n = 4, 6 and 6 mice. **e**, Validation of PES1 knockdown. Western blotting of PES1 in MCF10CA and SUM159 cells expressing control shRNA or PES1-targeting shRNAs (shRNA1 or shRNA2), with vinculin as a loading control. Bar graphs show PES1 protein levels normalized to vinculin and then to the control shRNA condition.

