## Supplementary Note 2 for "Ribosome biogenesis bottlenecks reveal vulnerabilities in cancer"

### Supplemental Note: Ribosome biogenesis bottlenecks reveal vulnerabilities in cancer

#### CONTENTS


I. Kinetic model for pre-rRNA production, processing, and degradation S2
   A. Model S2
   B. Loss function and fitting procedure S4
   C. Results for MCF10A and MCF10CA S4
   D. Results for inducible MYC overexpression S7
   E. Late-stage perturbations selectively reduce ribosome biogenesis efficiency S9

II. Spike-in measurements of RNA decay time S11

References S12

#### I. KINETIC MODEL FOR PRE-RRNA PRODUCTION, PROCESSING, AND DEGRADATION

##### A. Model

Here, we provide details of the mathematical model used in the main text to describe the dynamics of pre-ribosomal RNA (pre-rRNA) during the 5eU pulse-chase experiments (Fig. S1). In this model, 5eU is converted into intracellular 5eUTP at a rate $r_0$ and then incorporated into nascent rRNA transcripts at a rate $r_1$, which should be proportional to the transcription rate. The nascent rRNA ($s_0$) then undergoes a series of cleavage steps, each of which is described by a linear rate $k_i$, with $i$ labeling the site of cleavage. The cleavage of sites 01 and 02, which takes place quickly (confirmed by fitting, see Fig. S5A and Fig. S9A), is combined into one step with rate $k_{012}$ (green arrow). The cleavage of sites 1 and 2 can happen in any order (orange arrows, $k_1$ and $k_2$). For site 3', $\tilde{k}_{3'}$ denotes the rate of endonuclease cleavage at site 3' and $k_{3'}$ denotes the rate of gradual degradation by exonuclease until reaching ITS2-28S junction, which corresponds to the cleavage site measured by 5eU-seq. For site 4', steps 4a/b and 4' in Fig. S1A are simplified to a single step with rate $k_{4'}$ in the model (Fig. S1B). Each of the intermediates is also subject to degradation at a rate $k_d$ until it completes the last cleavage step, including sites 3, 3', or 4'. The intermediates allowed to degrade are enclosed by the dashed box.

The dynamics of these species are thus governed by the following set of ordinary differential equations (ODEs):

$$\frac{\mathrm{d}c_0}{\mathrm{d}t} = r_0\Theta(-t)\Theta(t+t_{\mathrm{pulse}}) - r_1 c_0, \tag{S1}$$

$$\frac{\mathrm{d}s_0}{\mathrm{d}t} = r_1 c_0 - (k_{012} + k_d)s_0, \tag{S2}$$

$$\frac{\mathrm{d}s_1}{\mathrm{d}t} = k_{012}s_0 - (k_1 + k_2 + k_d)s_1, \tag{S3}$$

$$\frac{\mathrm{d}s_2}{\mathrm{d}t} = k_2(s_1 + s_8) - (\tilde{k}_{3'} + k_d)s_2, \tag{S4}$$

$$\frac{\mathrm{d}s_3}{\mathrm{d}t} = \tilde{k}_{3'}s_2 - (k_{3'} + k_d)s_3, \tag{S5}$$

$$\frac{\mathrm{d}s_4}{\mathrm{d}t} = \tilde{k}_{3'}s_2 - (k_{4'} + k_d)s_4, \tag{S6}$$

$$\frac{\mathrm{d}s_5}{\mathrm{d}t} = k_2 s_1 - (k_1 + k_d)s_5, \tag{S7}$$

$$\frac{\mathrm{d}s_6}{\mathrm{d}t} = k_1 s_5 + k_2 s_8 - (k_3 + k_d)s_6, \tag{S8}$$

$$\frac{\mathrm{d}s_7}{\mathrm{d}t} = k_3 s_6, \tag{S9}$$

$$\frac{\mathrm{d}s_8}{\mathrm{d}t} = k_1 s_1 - (k_2 + k_d)s_8, \tag{S10}$$

$$\frac{\mathrm{d}s_9}{\mathrm{d}t} = k_{3'}s_3, \tag{S11}$$

$$\frac{\mathrm{d}s_{10}}{\mathrm{d}t} = k_{4'}s_4. \tag{S12}$$

For each of the post-cleavage rRNA species (18S, 5.8S, and 28S), the total concentration of all the precursors that contain these sequences is given by

$$S_{18\mathrm{S}} = s_0 + s_1 + s_5 + s_8 + s_6 + s_7, \tag{S13}$$

$$S_{5.8\mathrm{S}} = s_0 + s_1 + s_8 + s_2 + s_4 + s_{10}, \tag{S14}$$

$$S_{28\mathrm{S}} = s_0 + s_1 + s_8 + s_2 + s_3 + s_9. \tag{S15}$$

For each site ($i \in \{01, 02, 1, 2, 3, 3', 4'\}$), the fraction cleaved is defined as

$$f_i = \frac{c_i}{c_i + u_i}, \tag{S16}$$

where $c_i$ and $u_i$ are the amounts of cleaved and uncleaved pre-rRNA at site $i$, respectively. They are given by summing

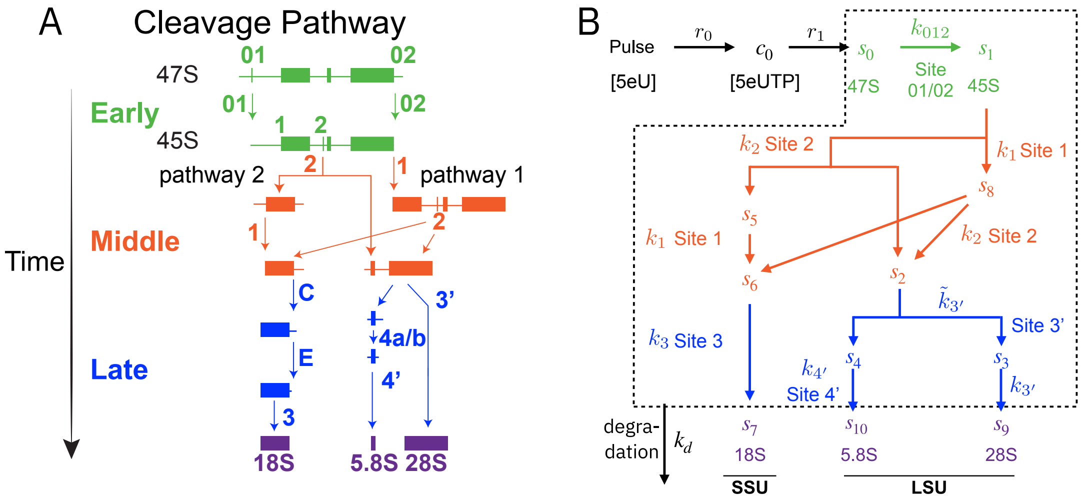


FIG. S1. Schematics for the rRNA cleavage pathway (A) and the mathematical model for rRNA production, processing, and degradation (B). In (B), "Site x" represents the cleavage at site x. In this model, 5eU is converted into intracellular 5eUTP at a rate $r_0$ and then incorporated into nascent rRNA transcripts at a rate $r_1$, which should be proportional to the transcription rate. The nascent rRNA ($s_0$) then undergoes a series of cleavage steps, each of which is described by a linear rate $k_i$, with $i$ labeling the site of cleavage. The cleavage of sites 01 and 02, which takes place quickly (confirmed by fitting, see Fig. S5A and Fig. S9A), is combined into one step with rate $k_{012}$ (green arrow). The cleavage of sites 1 and 2 can happen in any order (orange arrows, $k_1$ and $k_2$). For site 3', $\tilde{k}_{3'}$ denotes the rate of endonuclease cleavage at site 3' and $k_{3'}$ denotes the rate of gradual degradation by exonuclease until reaching ITS2-28S junction, which corresponds to the cleavage site measured by 5eU-seq. For site 4', steps 4a/b and 4' in Fig. S1A are simplified to a single step with rate $k_{4'}$ in the model (Fig. S1B). Each of the intermediates is also subject to degradation at a rate $k_d$ until it completes the last cleavage step, including sites 3, 3', or 4'. The intermediates allowed to degrade are enclosed by the dashed box. (A) is adapted from ref. [1].

over the relevant species that contain the cleaved or uncleaved sequence at site $i$:

$$u_{01} = s_0, \quad c_{01} = s_1 + s_5, \tag{S17}$$
$$u_{02} = s_0, \quad c_{02} = s_1 + s_8 + s_2 + s_3 + s_9, \tag{S18}$$
$$u_1 = s_0 + s_1 + s_5, \quad c_1 = s_6 + s_7 + s_8, \tag{S19}$$
$$u_2 = s_0 + s_1 + s_8, \quad c_2 = s_2 + s_4 + s_{10}, \tag{S20}$$
$$u_3 = s_0 + s_1 + s_5 + s_6, \quad c_3 = s_7, \tag{S21}$$
$$u_{3'} = s_0 + s_1 + s_8 + s_2 + s_3, \quad c_{3'} = s_9, \tag{S22}$$
$$u_{4'} = s_0 + s_1 + s_8 + s_2 + s_4, \quad c_{4'} = s_{10}. \tag{S23}$$

It is worth noting that the cleavage at downstream sites can affect the abundance of the upstream cleaved species. For example, after site 1 is cleaved, the sequence between sites 01 and 1 is degraded, which makes the cleavage of site 01 no longer detectable in the downstream intermediates.

The biogenesis efficiency (yield) can be defined for each cleavage site by quantifying the fraction of nascent pre-rRNA that undergoes cleavage at that site without being degraded. For each site ($i \in \{01, 02, 1, 2, 3', 4', 3\}$), the total flux that undergoes cleavage (i.e., cleaved before degraded) is given by

$$J_i = \int_{-t_{\text{pulse}}}^{\infty} j_i(t)\, \mathrm{d}t\,. \tag{S24}$$

The processing yield (efficiency) is defined as the fraction of transcripts that undergo cleavage without being degraded. It is calculated by taking the ratio of the total flux that undergoes cleavage versus that which is transcribed:

$$\eta_i = \frac{J_i}{J_0} = \frac{\int_{-t_{\text{pulse}}}^{\infty} j_i(t)\, \mathrm{d}t}{J_0}, \tag{S25}$$

where $J_0 = J_{\text{txn}} = r_0 t_{\text{pulse}}$ is the total amount of 5eU taken up and incorporated into pre-rRNA. The cleavage fluxes are defined as follows:

$$j_{01,02} = k_{012} s_0, \quad j_1 = k_1(s_1 + s_5), \quad j_2 = k_2(s_1 + s_8), \quad j_{3'} = k_{3'} s_3, \quad j_{4'} = k_{4'} s_4, \quad j_3 = k_3 s_6. \tag{S26}$$

In the main text and below, "amount of transcripts remaining" refers to the integrated flux that undergoes cleavage before degradation, $J_i$, and "fraction of transcripts remaining" refers to the processing efficiency, $\eta_i$.

##### B. Loss function and fitting procedure

Our model outputs four classes of quantities which can be measured experimentally: the fraction cleaved for each site ($f_i$), the spike-in normalized counts of cleaved and uncleaved pre-rRNA for each site ($c_i, u_i$), and the total counts of pre-rRNA that contain post-cleavage rRNA sequences ($S_{18\mathrm{S}}, S_{5.8\mathrm{S}}, S_{28\mathrm{S}}$). We infer the model parameters by maximum likelihood. For each class of measurements ($y \in \{\{f\}, \{c\}, \{u\}, \{S\}\}$), we can define the following negative log likelihood loss function to quantify the discrepancy between the model prediction $\hat{y}$ and the experimental measurement $y$:

$$\mathcal{L}_y = \frac{1}{2} \sum_{i\in\text{sites}} \sum_{t\in\text{time}} \left[ \frac{(y_i(t) - \hat{y}_i(t))^2}{\sigma_i^2(t) + \sigma_0^2} + \ln\left(\sigma_0^2 + \sigma_i^2(t)\right) \right], \tag{S27}$$

where $y_s(t)$ is the experimental measurement for site $s$ at time $t$ with estimated uncertainty $\sigma_s(t)$; $\hat{y}_s(t)$ is the model prediction; and $\sigma_0 \equiv \text{softplus}(\alpha_0) = \ln(1 + e^{\alpha_0})$ is an additional fitting parameter to account for potential underestimation of the experimental uncertainty[1]. Since the spike-in normalized counts ($y \in \{\{c\}, \{u\}, \{S\}\}$) are measured in arbitrary units, an additional scaling parameter is introduced for each of these measurements to rescale the experimental data $y_s(t) \to \beta_{y,s} y_s(t)$ before calculating the loss. No rescaling is needed for the fraction cleaved $\{f\}$.

The total loss function $\mathcal{L}$ is given by summing over many or all of the measurements. To examine whether the minimal model is able to describe the measured pre-rRNA dynamics, we only use the model to fit a subset of the measurements and use the rest as validation. For the results in the main text, we fit only $\{f\}$ and $\{S\}$ (i.e., $L_{\text{total}} = L_f + L_S$) and use $\{c\}$ and $\{u\}$ as validation. The kinetic parameters are parameterized as $k_i \equiv \text{softplus}(\theta_i)$ to ensure positivity. Fitting is done by minimizing the total loss function $L_{\text{total}}$ using the Adam optimizer [2] with respect to the transformed kinetic parameters $\theta_i$ and the additional uncertainty parameter $\alpha_0$.

The uncertainty of the fit is estimated using the Laplace approximation, where the covariance of the parameters $\theta_i$ is given by the inverse of the Hessian matrix of the loss function evaluated at the optimum. The uncertainty of the kinetic rates $k_i$ is then obtained by bootstrapping, which involves sampling the transformed parameters $\theta_i$ from a multivariate Gaussian distribution with the mean and covariance given by the Laplace approximation and then applying the softplus transformation to get $k_i$. The scaling parameters $\beta_{y,s}$ are optimized separately for each measurement and site by minimizing the loss function with respect to $\beta_{y,s}$ while keeping the kinetic parameters fixed, which is a least-squares problem with a closed-form solution. They are not included in the parameter uncertainty estimation. The uncertainty of all predicted quantities, such as ribosome biogenesis efficiency at each stage, is similarly estimated by bootstrapping, i.e., sampling the transformed kinetic parameters $\theta_i$ from a multivariate Gaussian distribution determined by the fit and measuring the mean and spread of the resulting predicted quantities.

##### C. Results for MCF10A and MCF10CA

Here, we present detailed results of fitting the model to the MCF10A and MCF10CA data shown in Fig. 3 of the main text. As described above, we split the experimental measurements into two sets, one for fitting and the other for validation. The fitting results are shown in Fig. S2, where the model could capture the dynamics of both the fraction cleaved and the total counts of pre-rRNA containing post-cleavage rRNA sequences for both MCF10A and MCF10CA cells. As validation, the model predicts the dynamics of the spike-in normalized counts of cleaved and uncleaved pre-rRNA for MCF10A (Fig. S3) and MCF10CA (Fig. S4), which were not used for fitting.

Fig. S5 shows the inferred kinetic parameters and ribosome biogenesis fluxes from the fitted model. The best fit cleavage rates (Fig. S5A) show comparable cleavage kinetics at early and middle sites (01, 02, 1, and 2) between MCF10A and MCF10CA, while the cleavage rates at late sites (3', 4', and 3) are significantly slower in MCF10CA than in MCF10A. This is consistent with the delay time analysis in Fig. 2 of the main text, although the timescales inferred here take into account concurrent transcription, cleavage, and degradation during the pulse-chase experiment, as well as the sequential cleavage order (Fig. S1). MCF10CA shows higher transcription activity than MCF10A (Fig. S5B).

---

[1] We use $\text{softplus}(x) = \ln(1 + e^x)$ as a smooth and differentiable way to parameterize positive parameters in the model, including the uncertainty floor parameter $\sigma_0$ and the rate parameters $k_i$.

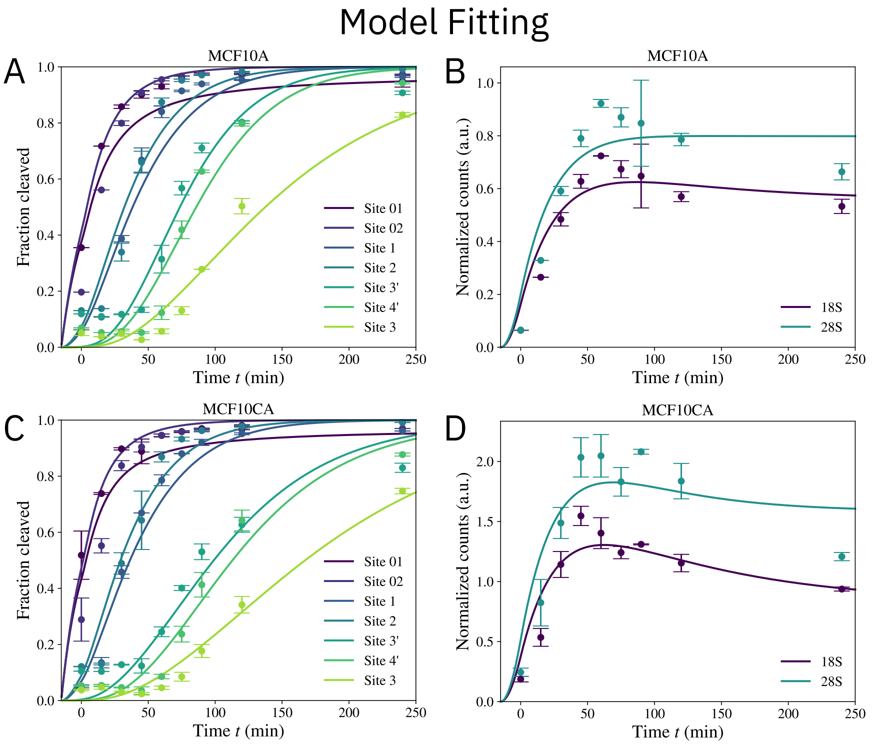


FIG. S2. Model fitting results for MCF10A and MCF10CA. The model was fitted to the fraction cleaved (A, C) and the total counts of pre-rRNA containing post-cleavage rRNA sequences (B, D). Data points are experimental measurements with error bars representing the standard deviation across replicates. Solid lines are model fits.

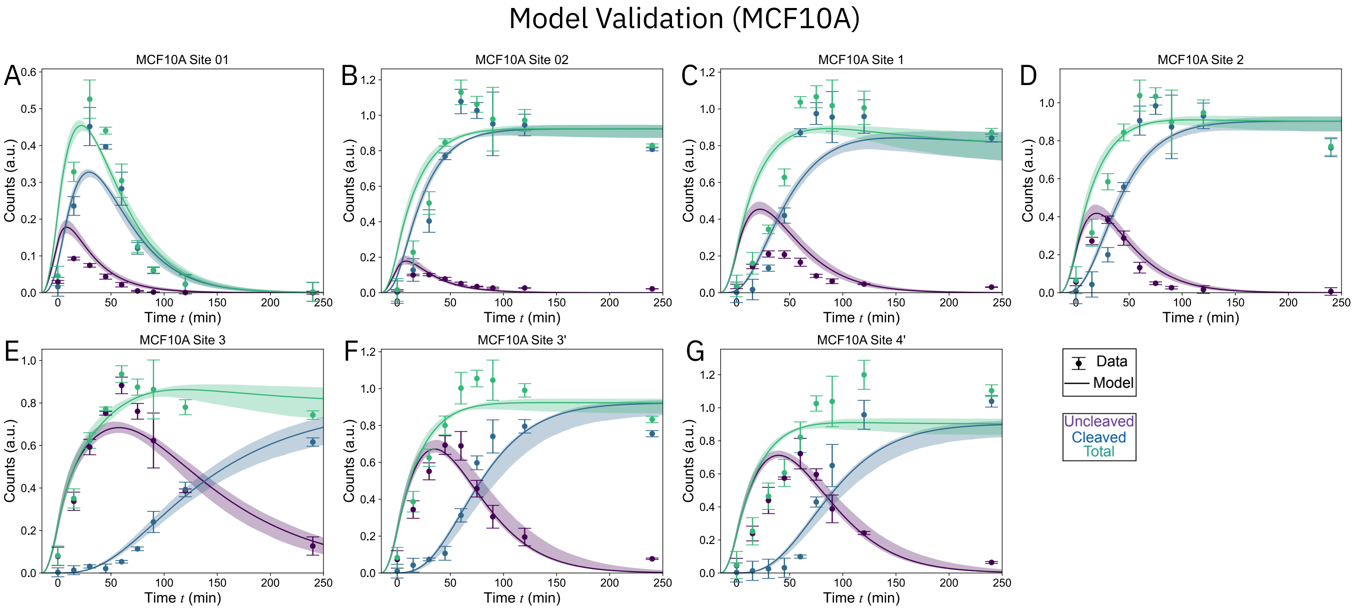


FIG. S3. Model validation for MCF10A. The model predictions for the spike-in normalized counts of cleaved and uncleaved pre-rRNA (solid lines) are compared with the experimental measurements (data points with error bars representing the standard deviation across replicates), which were not used for fitting. The shaded area represents model uncertainty estimated by bootstrapping. The experimental data were measured in arbitrary units and rescaled to match the model (see section I.B, *Loss function and fitting procedure* for details).

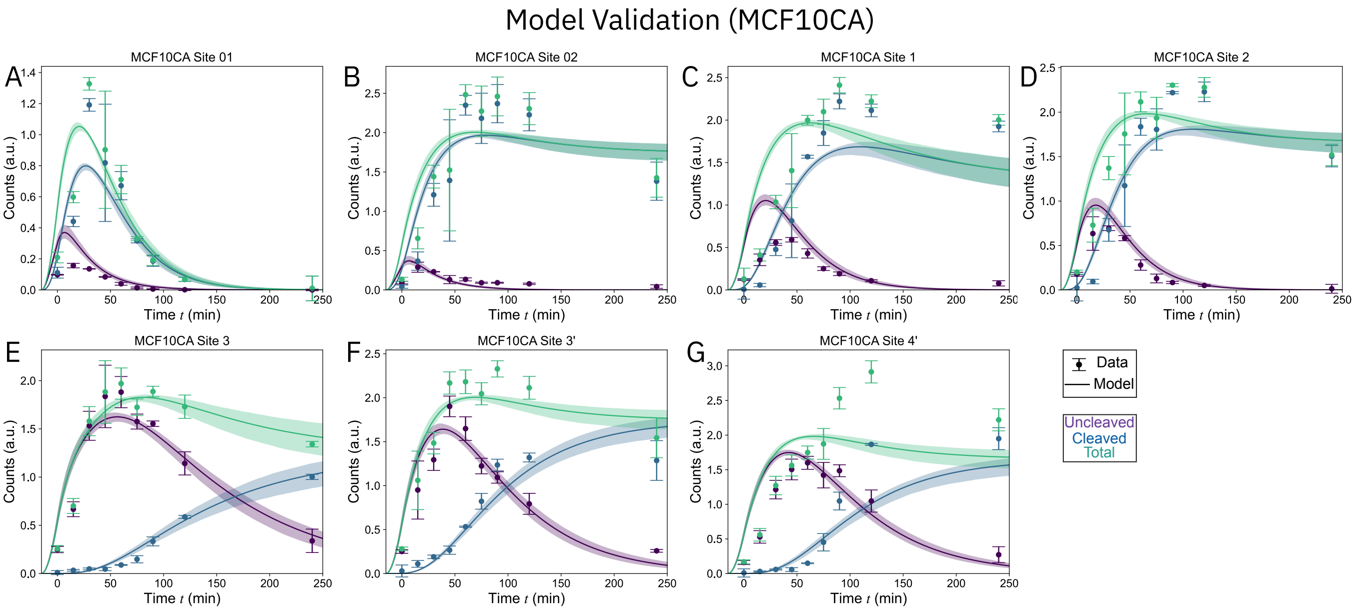


FIG. S4. Additional model validation results for MCF10CA. See Fig. S3 caption for a detailed description.

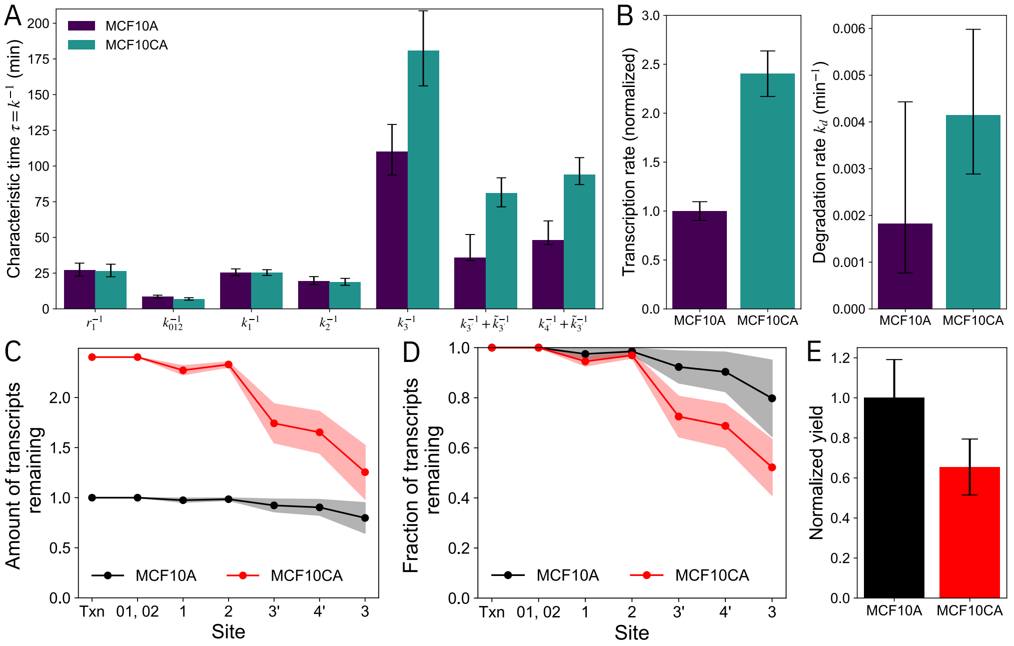


FIG. S5. Inferred pathway kinetic parameters and ribosome biogenesis fluxes from the fitted kinetic model. (A) Best fit kinetic parameters for MCF10A and MCF10CA, reported as characteristic timescales (i.e., inverse of the rates). Since 5eU-seq does not directly measure endonuclease cleavage at site 3', the inferred timescale for the experimentally measured cleavage site is given by the sum of the endonuclease cleavage time $\tilde{k}_{3'}^{-1}$ and the exonuclease degradation time $k_{3'}^{-1}$. For the same reason, the inferred timescale for site 4' is reported as the sum of $\tilde{k}_{3'}^{-1}$ and $k_{4'}^{-1}$, since the cleavage at site 4' also requires the endonuclease cleavage at site 3'. (B) Best fit transcription rate (as defined by the number of transcripts produced per time, normalized by MCF10A) and degradation rates $k_d$ for the pre-rRNA species that are allowed to degrade (i.e., those in the dashed box in Fig. S1B). (C) Amount of transcripts remaining, defined as the normalized flux for transcription (txn, $J_0$) and cleavage ($J_i$) at each site (see Eq. S24), normalized by the transcription activity ($J_0$) of MCF10A. (D) Fraction of transcripts remaining at each site, as defined by processing efficiency $\eta_i$ (Eq. S25). (E) Comparison of the normalized ribosome biogenesis efficiency, normalized by the efficiency of MCF10A (for easier comparison with Fig. 1e in the main text). In (A–B), the error bars represent the uncertainty estimated by the Laplace approximation. In (C–E), the error bars represent the uncertainty estimated by bootstrapping the model.

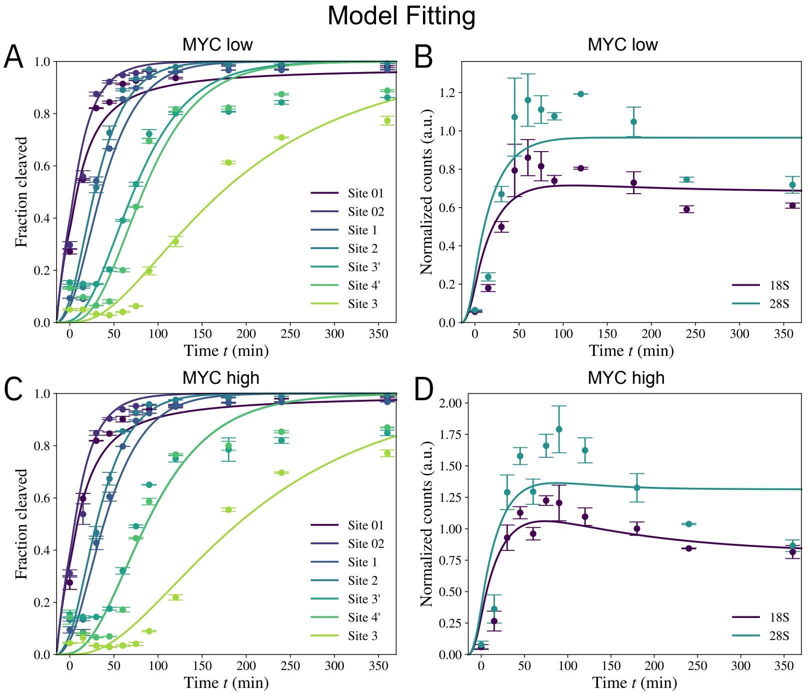


FIG. S6. Model fitting results for MYC-low and MYC-high cells. The model was fitted to the fraction cleaved (A, C) and the total counts of pre-rRNA containing post-cleavage rRNA sequences (B, D). Data points are experimental measurements with error bars representing the standard deviation across replicates. Solid lines are model fits.

Both cell lines show non-negligible degradation (Fig. S5B), which is required for explaining the decay of post-cleavage-containing pre-rRNA species at long chase times (Fig. S2B, D). Interestingly, the degradation rate appears slightly higher in MCF10CA, although the difference cannot be fully resolved given the uncertainty of the fit.

From the model, we can infer the flux of pre-rRNA that undergoes cleavage at each site (Eq. S24 and Fig. S5C). Consistent with the measurements in Fig. 1 of the main text for the inducible MYC overexpression system, we find that oncogenic cells (MCF10CA) have a higher transcription activity (i.e., higher $J_0$, see "Txn" in Fig. S5C) as well as higher cleavage flux at late sites (3', 4', and 3), which are indicative of increased ribosome production. The processing efficiency (Eq. S25 and Fig. S5D) maintains close to 100% for early and middle sites (01, 02, 1, and 2) in both cell lines. It is reduced in late sites (3', 4', and 3) in both cell lines, with a more significant reduction in MCF10CA than in MCF10A. To estimate the final efficiency, we use the cleavage of site 3, since it takes place after cytoplasmic export and is thus more indicative of the amount of mature ribosomes produced. By contrast, sites 3' and 4' are cleaved in the nucleus and followed by additional processing steps and cytoplasmic export, which may involve additional reduction in efficiency. By this measure, we estimate the normalized ribosome biogenesis efficiency to be $80 \pm 15\%$ for normal (MCF10A) cells and $52 \pm 11\%$ for malignant (MCF10CA) cells. Normalizing the efficiency (yield) by MCF10A (Fig. S5E) for MCF10A, we find a relative reduction of $35 \pm 10\%$ in MCF10CA compared to MCF10A. This is comparable if not slightly higher than the relative efficiency difference measured for the inducible MYC overexpression system (see Fig. 1e in the main text), although direct experimental measurement of ribosome biogenesis efficiency using endogenous ribosomal protein tagging was not performed in MCF10CA cells because clonal selection required for this assay consistently altered cellular fitness and deviated from the parental malignant phenotype (Supplementary Note 4).

##### D. Results for inducible MYC overexpression

Next, we analyze the inducible MYC overexpression system (Fig. 1 and Fig. 3f of the main text) using the same procedure described above for MCF10A and MCF10CA. Here, we pool together data from two separate pulse-chase experiments, performed for 0–4 hours chase time and 1–6 hours chase time, respectively. The data were aligned at 2 hours.

The model was fit to the fraction cleaved and the total counts of pre-rRNA containing post-cleavage rRNA sequences

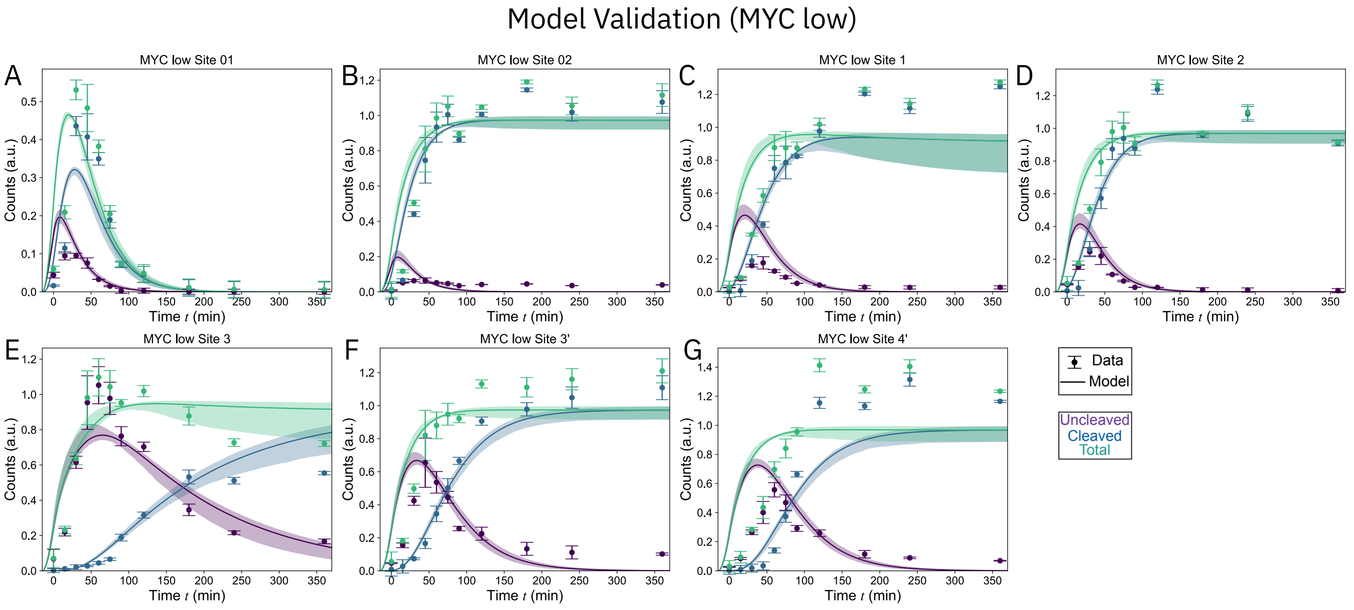


FIG. S7. Model validation results for MYC-low. See Fig. S3 caption for a detailed description.

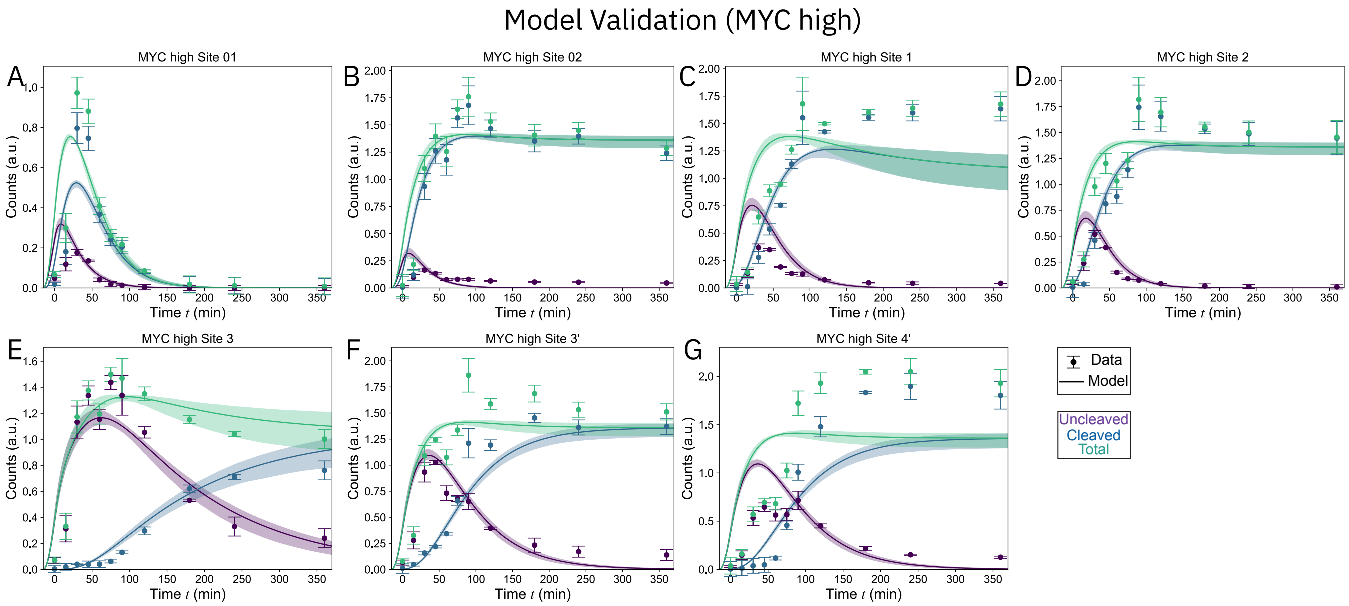


FIG. S8. Model validation results for MYC-high. See Fig. S3 caption for a detailed description.

for MYC-low and MYC-high cells, with the results shown in Fig. S6. Validation using the spike-in normalized counts of cleaved and uncleaved pre-rRNA was shown in Fig. S7 and Fig. S8. The inferred kinetic parameters (Fig. S9A) reveal comparable cleavage kinetics at early and middle sites (01, 02, 1, and 2) between MYC-low and MYC-high cells, while the cleavage rates at late sites (3', 4', and 3) are significantly slower in MYC-high cells than in MYC-low cells. MYC-high shows a $\sim 50\%$ increase in transcription activity (Fig. S9B), consistent with measurement by 5eU-imaging (Fig. 1d) in the main text. The degradation rate appears slightly higher in MYC-high, although the difference cannot be fully resolved given the uncertainty of the fit.

Computing the total cleavage flux $J_i$ (i.e., the amount of transcripts remaining at a given cleavage step, see Fig. S9C) and the processing efficiency (i.e., the fraction of transcripts remaining, see Fig. S9D) as defined above, we find that most of the efficiency reduction takes place during late-stage cleavage. Accordingly, the fraction of nascent pre-rRNA completing the final cytoplasmic cleavage step (site 3) decreased from $90 \pm 14\%$ in MYC-low cells to $67 \pm 14\%$ in

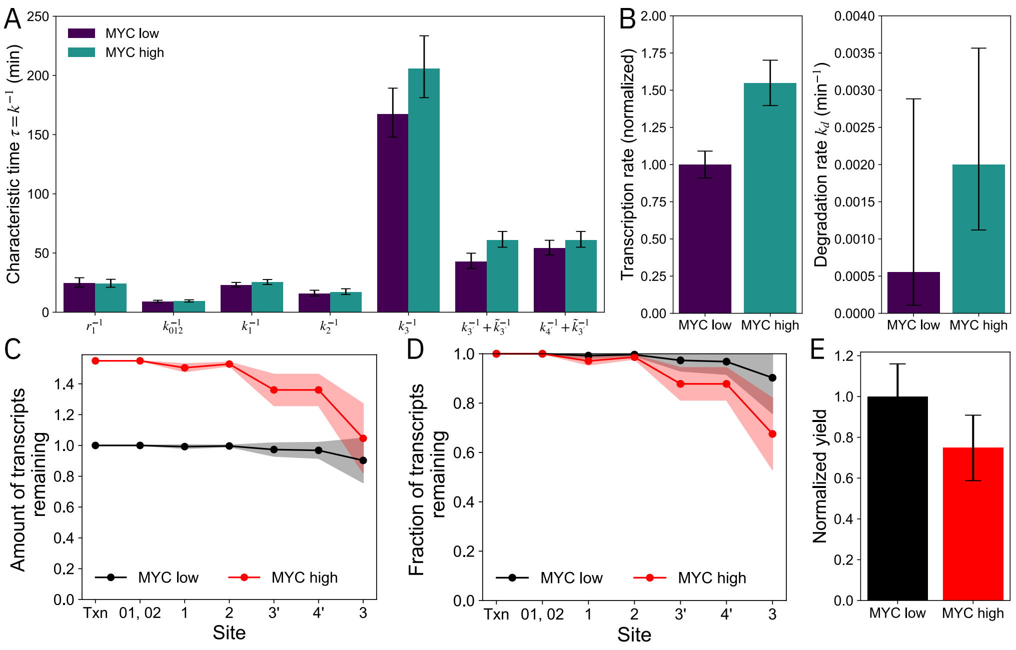


FIG. S9. Inferred pathway kinetic parameters and ribosome biogenesis fluxes from the fitted kinetic model, for MYC-low and MYC-high cells. See Fig. S5 caption for a detailed description of the plotted quantities.

MYC-high cells. To compare the model-derived estimate with experimental measurements, we normalize the efficiency by that of MYC-low, which reveals a $25\% \pm 16\%$ relative reduction in efficiency in MYC-high cells (Fig. S9E). This is fully consistent with independent measurements using 5eU labeling of nascent rRNA transcription and TMR labeling of newly synthesized ribosomal subunits, which revealed a similar decrease in efficiency ($29 \pm 10\%$ for LSU via HaloTag-RPL10A and $25 \pm 10\%$ for SSU via HaloTag-RPS14) (Extended Data Fig. 1e–l and Fig. 1e).

##### E. Late-stage perturbations selectively reduce ribosome biogenesis efficiency

In the model, we can mimic the effect of knocking down processing factors by reducing the corresponding cleavage rates. If late-stage processing forms a bottleneck that leads to reduced ribosome biogenesis efficiency in oncogenic cells, we would expect that perturbing the cleavage rates at late sites (3', 4', and 3) would lead to a more significant reduction in the ribosome biogenesis efficiency in MYC-high cells than in MYC-low cells. To illustrate this conceptual point, we assume the simplest case where the cleavage rates decrease by the same factor in MYC-low and MYC-high cells. The model-predicted ribosome biogenesis efficiency is shown in Fig. S10, for perturbing the cleavage rates at site 3 (Fig. S10A) and site 3' (Fig. S10B). The efficiency is normalized by the efficiency of the unperturbed MYC-low cells. As the cleavage rate decreases, the efficiency decreases in both MYC-low and MYC-high cells (from top right to bottom left), but the reduction is more significant in MYC-high cells than in MYC-low cells, as demonstrated by the convex shape of the model prediction (solid black lines) in Fig. S10A and B. Similar results were found in the best-fit model for MCF10A and MCF10CA cells (Fig. S10C and D).

Consistent with the theoretical prediction, the experimentally measured changes in ribosome biogenesis efficiency upon knockdown of late processing factors, including PES1, which is required for ITS2 processing during large subunit (LSU) maturation, and NOB1, the cytoplasmic endonuclease that performs the final cleavage at site 3 to generate mature 18S rRNA (Extended Data Fig. 5b–e). All measured biogenesis efficiencies fall on the theory lines (Fig. S10A–B).

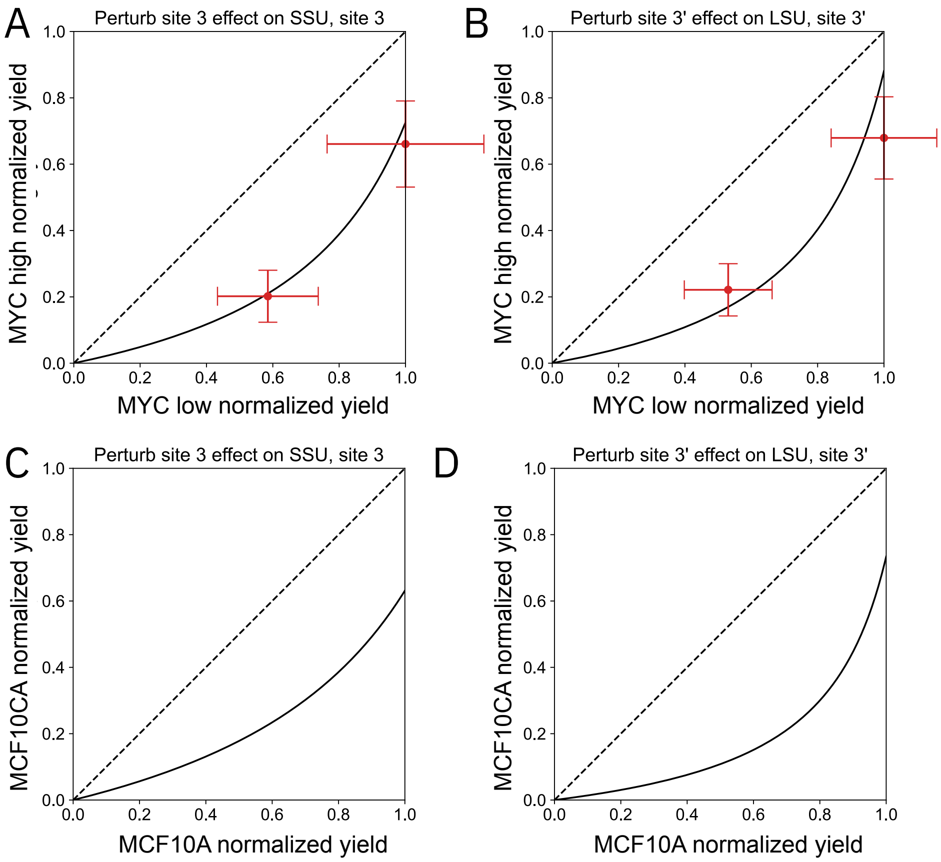


FIG. S10. Model prediction for the effect of perturbing the cleavage rates at site 3 or site 3' on the ribosome biogenesis efficiency in MYC-low/high cells (A–B) and MCF10A/MCF10CA cells (C–D). The efficiency is normalized by the efficiency of the unperturbed MYC-low cells (A–B) or the unperturbed MCF10A cells (C–D). In all panels, the solid black lines are the model prediction: (A, C) Perturbation of site 3 by reducing $k_3$. (B, D) Perturbation of site 3 by reducing $\tilde{k}_{3'}$. Data points and error bars represent experiments for NOB1 knockdown (A) and PES1 knockdown (B) (see Extended Data Fig. 5b–e).

#### II. SPIKE-IN MEASUREMENTS OF RNA DECAY TIME

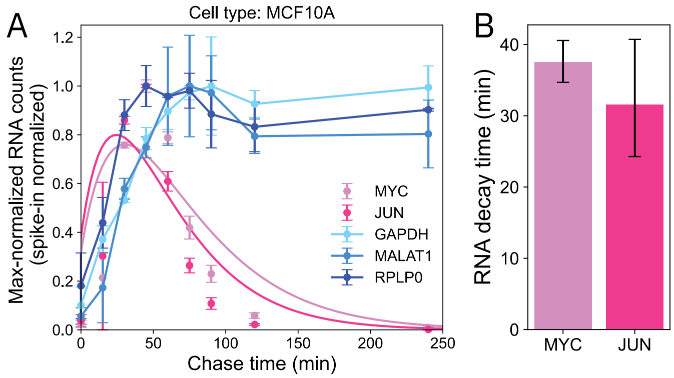


FIG. S11. Characterization of RNA degradation using spike-in normalized 5eU-seq. (A) Spike-in normalized RNA counts. Data points are experiments with error bars representing the standard deviation across replicates. For short-lived RNA (red), solid lines are fits to the model in Eq. S31 with the inferred degradation time shown in (B). For long-lived RNA (blue), the solid line simply connects the data points to guide the eye.

In this section, we analyze the dynamics of several non-ribosomal RNA during the 5eU pulse-chase experiment to demonstrate that the spike-in-normalized RNA abundance can be used to quantify RNA degradation over time. To describe the production and degradation dynamics of the RNAs, we consider the following model:

$$\frac{\mathrm{d}c}{\mathrm{d}t} = r_0 \Theta(-t)\Theta(t + t_{\text{pulse}}) - r_1 c, \tag{S28}$$

$$\frac{\mathrm{d}s}{\mathrm{d}t} = r_1 c - k_d s, \tag{S29}$$

where $c$ and $s$ denote the abundance of 5eUTP and RNA, respectively; $r_0$ is the rate of converting 5eU to 5eUTP, $r_1$ is the rate of eU incorporation via transcription, and $k_d$ is the RNA degradation rate. In the first equation, the factor $\Theta(-t)\Theta(t + t_{\text{pulse}})$ indicates that 5eU is only present during the incubation period (pulse) $t \in (-t_{\text{pulse}}, 0)$.

The solutions are

$$c(t) = \frac{r_0}{r_1} \begin{cases} 1 - e^{-r_1(t+t_{\text{pulse}})} & t \in (-t_{\text{pulse}}, 0) \\ (1 - e^{-r_1 t_{\text{pulse}}})e^{-r_1 t} & t \in (0, +\infty) \end{cases}, \tag{S30}$$

$$s(t) = \begin{cases} \frac{r_0}{k_d}\left(1 - e^{-k_d(t+t_{\text{pulse}})}\right) + \frac{r_0}{k_1 - k_d}\left(e^{-k_1(t+t_{\text{pulse}})} - e^{-k_d(t+t_{\text{pulse}})}\right) & t \in (-t_{\text{pulse}}, 0) \\ s(0)e^{-k_d t} + \frac{r_0(1-e^{-r_1 t_{\text{pulse}}})}{r_1 - k_d}\left(e^{-k_d t} - e^{-r_1 t}\right) & t \in (0, +\infty) \end{cases}, \tag{S31}$$

where $s(0) = \frac{r_0}{k_d}\left(1 - e^{-k_d t_{\text{pulse}}}\right) + \frac{r_0}{r_1 - k_d}\left(e^{-r_1 t_{\text{pulse}}} - e^{-k_d t_{\text{pulse}}}\right)$.

Thus, for each RNA species, we can determine the degradation rate $k_d$ by fitting the spike-in normalized RNA counts to $s(t)$ $(t > 0)$, up to a scaling factor that can be absorbed into $r_0$. As shown in Fig. S11, the model could quantify the degradation time of relatively short-lived RNA (in red), while the lifetime of long-lived RNA (in blue) could not be determined since it is longer than the duration of the experiment. The decay times are consistent with previous measurements [3–5].
